# Psilocybin reshapes cortical inhibition through selective interneuron recruitment

**DOI:** 10.64898/2026.04.16.718963

**Authors:** Pasha A. Davoudian, Quan Jiang, Cory A. Knox, Neil K. Savalia, Ling-Xiao Shao, Joshua Wilson, Amanda M. Weiner, Catherine W. Chong, Clara Liao, Jack D. Nothnagel, Takeshi Sakurai, Alex C. Kwan

## Abstract

Psychedelics show therapeutic potential for treating psychiatric disorders. While studies have emphasized the roles of cortical pyramidal cells, GABAergic neurons also express serotonin receptors and are therefore likely targets of psychedelics. In this study, we determine the effect of psilocybin on the activity dynamics of major GABAergic cell types in the mouse medial frontal cortex. Psilocybin reduces the firing of somatostatin-expressing interneurons, but increases the activity of parvalbumin-expressing interneurons. This cell type-specific response is unlikely to involve vasoactive intestinal peptide-expressing interneurons. Instead, pharmacological blockade and conditional knockout experiments demonstrate that psilocybin acts on the 5-HT_1A_ receptor at SST interneurons, which contributes to the drug’s long-term behavioral effects. Collectively, the results reveal that the classic psychedelic psilocybin alters cortical inhibition in a cell type-specific manner.

## MAIN TEXT

Psilocybin is a classic psychedelic that may be a promising therapeutic for people living with various psychiatric conditions (*1, 2*). Clinical trials have demonstrated that just one or two sessions of psilocybin-assisted therapy can rapidly reduce symptoms of depression and maintain this remission for several weeks to months (*3–7*). The potential therapeutic value underscores a need to understand the neurobiology underlying psilocybin’s drug action.

Most studies of psychedelics in the neocortex have focused on the excitatory pyramidal cells. However, this emphasis is not fully justified, because classic psychedelics are partial or full agonists at numerous serotonin receptors, including the 5-HT_2A_, 5-HT_1A_, and 5-HT_2C_ subtypes that are expressed in cortical inhibitory neurons (*8*). Indeed, serotonin modulates inhibitory synaptic currents and electrophysiological properties of cortical interneurons in brain slices as well as *in vivo* (*9–12*). For psychedelics, several studies tested the effects of 2,5-dimethoxy-4-iodoamphetamine (DOI) on interneurons in brain slices (*13, 14*) and on cortical oscillations (*15–17*). Recent transcriptomics studies hinted at psychedelics’ effects on cortical interneuron populations at the molecular level (*8, 18*). These studies suggest psychedelic drug actions on GABAergic neurons, yet we lack a clear picture of how classic psychedelics influence the firing dynamics of interneurons *in vivo*, particularly for compounds like psilocybin that are being evaluated for clinical use.

The impact of psychedelics on GABAergic neurons in the neocortex is likely to depend on cell type. One way to classify cortical interneurons is based on the expression of parvalbumin (PV), somatostatin (SST), or vasoactive intestinal peptide (VIP), which together make up two thirds of the GABAergic cells in the mouse frontal cortex (*19*). These GABAergic cell types have distinct connectivity: PV and SST interneurons target the somatic and dendritic compartments of pyramidal cells respectively to regulate the activity of pyramidal cells (*20, 21*). Wiring among interneurons is stereotyped, with PV interneurons strongly inhibiting other PV interneurons (*22–24*), SST interneurons inhibiting most other neurons (*22, 23*), and VIP interneurons inhibiting SST interneurons in a disinhibitory circuit motif (*22, 25, 26*). Functionally, cortical PV, SST, and VIP interneurons exhibit distinct patterns of firing activity during various forms of active behavior (*27–32*). Clarifying the effect of psychedelics on the activity of interneuron populations will be an essential step towards understanding how the drugs shape cortical activity and modify behavior.

In this study, we use cell type-specific extracellular electrophysiology and two-photon calcium imaging to measure the activity of GABAergic neurons in the mouse medial frontal cortex *in vivo*. We find that psilocybin acutely decreases the firing of SST interneurons but increases spiking of PV interneurons. The suppressive effect of psilocybin on SST interneurons is unlikely to involve the disinhibitory circuit motif, because we did not detect any psilocybin-induced activity change in VIP interneurons. Rather, 5-HT_1A_ receptor expression in SST interneurons is a key target of psilocybin’s effects, as supported by pharmacological blockade and conditional knockout experiments. Deletion of the 5-HT_1A_ receptor in SST interneurons interferes with psilocybin’s long-term beneficial effects on stress-related behaviors. Together, the results demonstrate psilocybin’s opposing effects on the firing of the major interneuron populations in the neocortex and pinpoint a 5-HT_1A_ receptor-dependent mechanism.

### Psilocybin suppresses the firing of frontal cortical SST interneurons

We recorded spontaneous spiking activity from genetically identified SST interneurons in the medial frontal cortex of awake, head-fixed mice. We used *Sst^Cre^*;Ai32 mice, which were generated by crossing a *Sst^Cre^* mouse (*33*) with an Ai32 reporter mouse (*34*), resulting in Cre-dependent expression of ChR2 in SST interneurons. We performed a craniotomy in the mouse frontal cortex above the ACAd and medial MOs region of the medial frontal cortex and inserted a high-density Neuropixels 1.0 electrode (*35*) into the brain (**Figure 1A**). We recorded for 30 minutes, injected psilocybin (1 mg/kg, i.p.) or saline via a catheter, and then recorded for another 60 minutes (**Figure 1B**). The catheter allowed for drug delivery without needle insertion during the recording, which could agitate the animal to cause motion artifacts. At the end of each recording session, we performed “opto-tagging”, where we used an optical fiber to deliver trains of brief laser pulses (20-50 ms, 473 nm) onto the cortical surface to elicit spiking in the ChR2-expressing cells. Animals would be sacrificed after the recording for histology. The Neuropixels probe was coated with the fluorescent dye CM-DiI, allowing it to be visualized in fixed brain sections post hoc to confirm placement in the correct brain region (**Figure 1C and 1D**).

**Figure 1:**
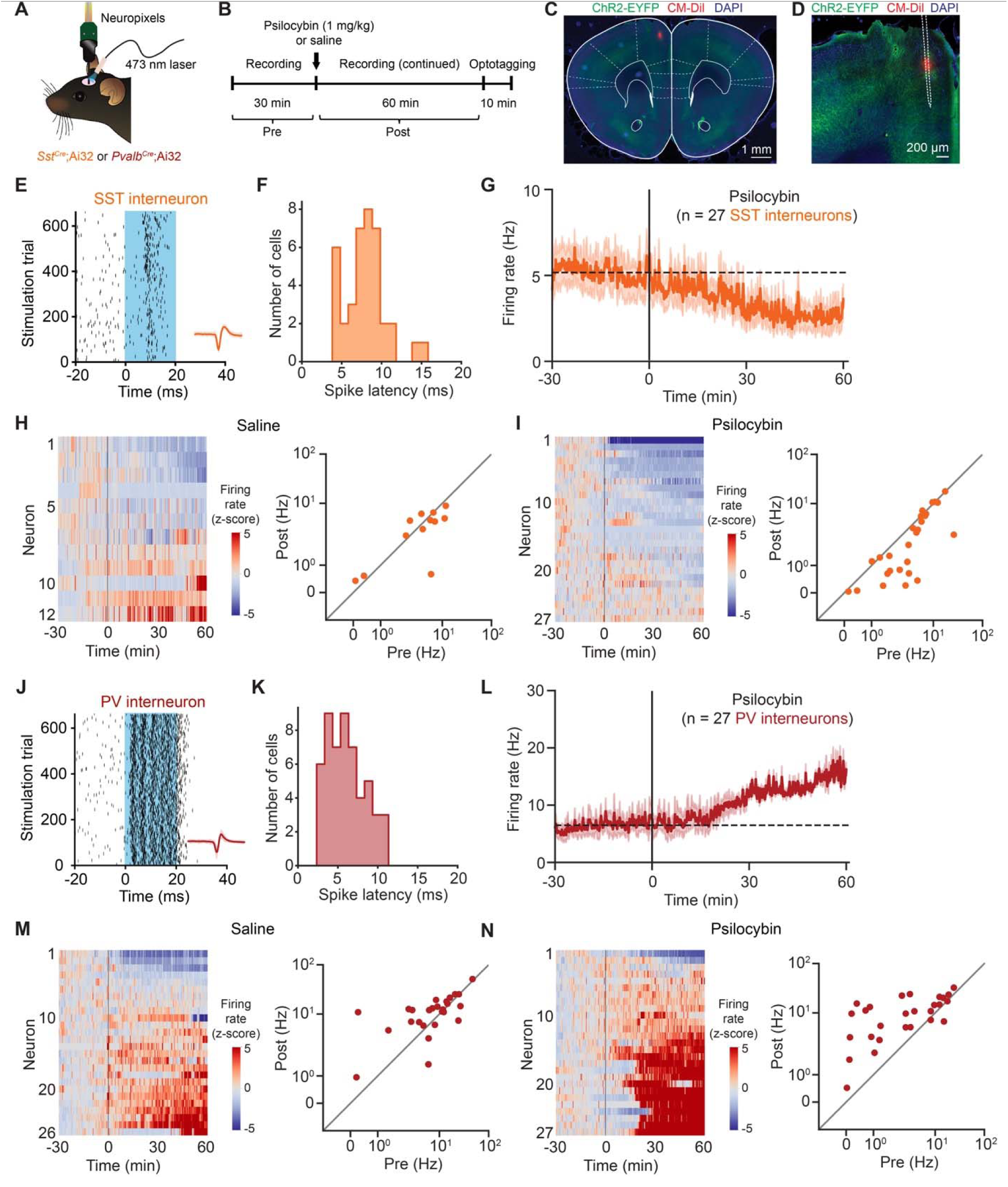
Psilocybin evokes distinct firing changes in SST and PV interneurons in the mouse medial frontal cortex. **(A)** Spiking activity of ChR2-expressing SST interneurons in *Sst^Cre^*;Ai32 mice or PV interneurons in *Pvalb^Cre^*;Ai32 mice was recorded using a Neuropixels probe. **(B)** The experimental timeline includes pre-drug (−30 – 0 min) and post-drug (0 – 60 min) periods. **(C)** Coronal histological section, showing the targeting of the Neuropixels probe that was coated with CM-DiI dye (red), along with ChR2-GFP expression (green) and cell nuclei stained via DAPI (blue). **(D)** Magnified view of (C). **(E)** Spike raster of an opto-tagged SST interneuron from a *Sst^Cre^*;Ai32 mouse. Blue, period of laser stimulation. Inset, the spike waveform, mean ± s.d. **(F)** The latency to the first spike after the onset of laser stimulation for opto-tagged SST interneurons. n = 39 cells from 12 mice. **(G)** Firing rate before and after psilocybin administration, averaged across opto-tagged SST interneurons. Line, mean. Shading, ± SEM. Dashed line, mean firing rate during pre-drug period. n = 27 cells from 9 mice. **(H)** Left, the activity of opto-tagged SST interneurons before and after saline administration. The firing rate of each neuron was converted to a z-score by normalization based on its pre-drug firing rate. Right, mean firing rates during pre- and post-drug periods for the opto-tagged SST interneurons. The axes use a symmetric logarithmic scale, which includes a linear region (0 to 0.03 Hz) that transitions to a logarithmic region (>0.03 Hz). Each dot represents one neuron. n = 12 cells from 3 mice. **(I)** Similar to (H) for psilocybin administration. n = 27 cells from 9 mice. **(J - N)** Similar to (E - I) for opto-tagged PV interneurons from *Pvalb^Cre^*;Ai32 mice. n = 26 cells from 3 mice for saline, and 27 cells from 3 mice for psilocybin.

We recorded from 12 *Sst^Cre^*;Ai32 animals: 9 mice were given psilocybin (4 males, 5 females) and 3 mice were given saline (1 male, 2 females). We isolated single units by automated spike sorting, manual curation, and then included those that passed four quality metrics: presence ratio, inter-spike-interval violations ratio, amplitude cutoff, and isolation distance (**Figure S1**). For psilocybin, we had 1021 cells including 27 opto-tagged SST interneurons (range: 82–330 cells and 1–5 opto-tagged units per animal). For saline, we had 212 cells including 12 opto-tagged SST interneurons (range: 65–77 cells and 1–8 opto-tagged units per animal). **Figure 1E** shows the spike raster of an opto-tagged SST interneuron, which was driven to spike by the laser photostimulation. Across all opto-tagged SST interneurons, latency from laser onset to first spike was 7.9±0.4 ms (mean ± SEM; **Figure 1F**) and the baseline firing rate during pre-drug period was 5.4±0.8 Hz (mean ± SEM).

After the administration of psilocybin, SST interneurons on average reduced their spiking activity (**Figure 1G**). Comparing their firing rates between the pre-drug (−30 to 0 minute) and post-drug period (0 to 60 minute), there was no change after saline injection (pre-saline: 5.0±1.1Hz, post-saline: 3.9±0.8 Hz; **Figure 1H, S2, S3**), but there was a significant decrease in spike rate after psilocybin administration (pre-psilocybin: 5.5±1.1 Hz, post-psilocybin: 3.7±0.8 Hz; **Figure 1I, S2, S3**). Some cells were recorded from the same mouse, therefore we fit the drug-evoked change in firing rate for SST interneurons using a mixed effects model, which accounted for the hierarchical structure of the data (main effect of treatment: *P* = 0.028, mixed effects model; see **Methods**). If we consider a mean z-score of −2 during the post-drug period to be a substantial decrease in spike rate, then 18.5% of the recorded SST interneurons reduced firing activity following psilocybin administration, whereas none of the recorded SST interneurons exhibited such change after saline (psilocybin: 5 cells with Z < −2, 0 cell with Z > 2, out of 27 SST interneurons; saline: 0 cell with Z < −2, 1 cell with Z > 2, out of 12 SST interneurons). These results show that psilocybin administration led to an acute reduction in the spiking activity of SST interneurons in the mouse medial frontal cortex.

### Psilocybin increases the firing of frontal cortical PV interneurons

In addition to SST interneurons which constitute ∼20% of GABAergic cells in the mouse frontal cortex, PV interneurons are the other major population that accounts for ∼40% of the inhibitory interneurons (*19*). We therefore next asked how psilocybin affects the spiking activity of PV interneurons. We generated *Pvalb^Cre^*;Ai32 mice by crossing a *Pvalb^Cre^* mouse (*36*) with a Ai32 reporter mouse for Cre-dependent expression of ChR2 in PV interneurons. We used the same Neuropixels and opto-tagging approach, recording before and after the administration of psilocybin (1 mg/kg, i.p.) or saline. We recorded from 6 *Pvalb^Cre^*;Ai32 animals: 3 mice were given psilocybin (2 males, 1 female) and 3 mice were given saline (1 male, 2 females). For psilocybin, we had 338 cells including 27 opto-tagged PV interneurons (range: 99–127 cells and 5–11 opto-tagged units per animal). For saline, we had 504 cells including 26 opto-tagged PV interneurons (range: 106–205 cells and 4–17 opto-tagged units per animal; **Figure S1**). Across all opto-tagged PV interneurons, the latency from laser onset to first spike was 6.0±0.3 ms (mean ± SEM; **Figure 1J and 1K**) and the baseline firing rate during pre-drug period was 8.6±1.3 Hz (mean ± SEM).

In contrast to what we observed in SST interneurons, the PV interneurons had an increase in firing rate that occurred ∼15 minutes after psilocybin injection (**Figure 1L**). Comparing the pre-and post-drug firing rates revealed a significant elevation of spike rate after the administration of psilocybin (pre-saline: 11.3±2.1 Hz, post-saline: 13.2±2.0 Hz; pre-psilocybin: 6.1±1.4 Hz, post-psilocybin: 11.6±1.6 Hz; main effect of treatment: *P* = 0.062, mixed effects model; **Figure 1M and 1N, S4, S5**). About half of the recorded PV interneurons had a substantial rise in spiking activity after psilocybin administration (psilocybin: 14 cells with Z > 2, 1 cell with Z < −2, out of 27 PV interneurons). Based on probe reconstruction, the majority of recorded SST and PV interneurons were located in layer 5 (**Figure S4**). Therefore, psilocybin has opposing effects on the spike rate of the two major populations of GABAergic neurons in the medial frontal cortex.

### VIP interneurons are unlikely to mediate psilocybin’s suppressive effect on SST interneurons

We were particularly interested in psilocybin’s suppressive effect on the SST interneurons. SST interneurons target the dendritic compartments of pyramidal cells, so a reduction in their activity should disinhibit dendrites. This can result in a higher level of dendritic electrogenesis, potentially enabling psilocybin’s effect on structural neural plasticity (*37*). To gain insight into the potential mechanism, we analyzed the publicly available single-cell sequencing data set from the Allen Institute for Brain Science (*38*). We examined transcripts encoding serotonin receptor subtypes, *Htr1a*, *Htr2a*, *Htr2b*, *Htr2c*, and *Htr3a*, in VIP, SST, and PV interneurons (**Figure 2A**). The expression of serotonin receptor transcripts in the mouse frontal cortex has cell type-specific preference. VIP interneurons predominantly express *Htr2c* (65% of cells, relative to 6% and 8% for *Htr1a* and *Htr2a*). SST interneurons express *Htr1a* more than other subtypes (39% of cells, relative to 18% and 15% for *Htr2a* and *Htr2c*). PV interneurons primarily express *Htr2a* (47% of cells, relative to 4% and 35% for *Htr1a* and *Htr2c*). VIP interneurons additionally express *Htr3a*, but should not contribute to drug action because psilocin has negligible binding to this ionotropic serotonin receptor (*39*). *Htr2c* encodes the 5-HT_2C_ receptor, thought to be a G_q_-coupled receptor that increases neuronal excitability. By contrast, *Htr1a* encodes the 5-HT_1A_ receptor, which is a G_i_-coupled receptor known to decrease neuronal excitability (*40, 41*). Thus, there are two plausible mechanisms for psilocybin’s suppressive effects on SST interneurons. One, psilocybin may act via 5-HT_2C_ receptors at VIP interneurons, which would increase their firing, leading to inhibition of SST interneurons. Two, psilocybin may act via 5-HT_1A_ receptors at SST interneurons to inhibit directly their firing. These mechanisms are not mutually exclusive.

**Figure 2:**
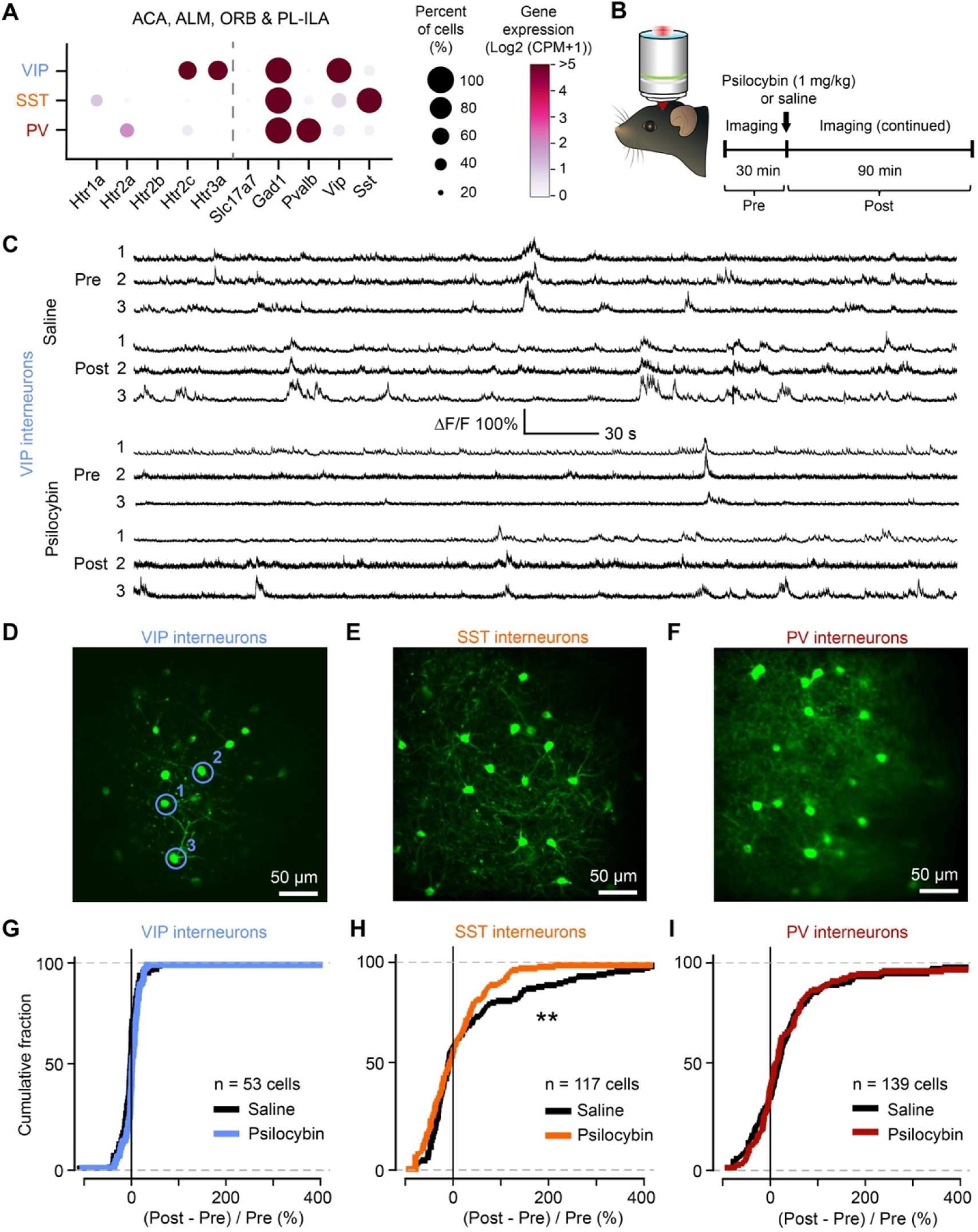
Psilocybin did not change the activity of VIP interneurons in the mouse medial frontal cortex. **(A)** Single-cell transcript counts summed from the mouse ACA, ALM, ORB, and PL-ILA regions, extracted from the SMART-Seq data set from Allen Institute. Percent of cells denote the fraction that has non-zero number of transcripts. *Slc17a7* and *Gad1* are markers of excitatory and GABAergic neurons, respectively. *Vip, Sst,* and *Pvalb* are markers of GABAergic cell populations. ACA, anterior cingulate area; ALM, anterior-lateral motor area; ORB, orbital area; PL, prelimbic area; ILA, infra-limbic area. **(B)** Head-fixed mice were imaged using a two-photon microscope. The experimental timeline includes pre-drug (−30 – 0 min) and post-drug (0 – 90 min) periods. **(C)** Spontaneous calcium transients from cell bodies of GCaMP6f-expressing VIP interneurons in the medial frontal cortex of a *Vip^Cre^* mouse. Each row is a cell. The same three cells are shown for a 5-minute epoch within the pre- and post-drug periods for saline and pre- and post-drug periods for psilocybin. **(D)** GCaMP6f-expressing VIP interneurons in the field of view sampled during *in vivo* two-photon imaging. Circles, neurons corresponding to the transients shown in (C). **(E)** Similar to (D) for SST interneurons in a *Sst^Cre^* mouse. **(F)** Similar to (D) for PV interneurons in a *Pvalb^Cre^* mouse. **(G)** Change in the rate of spontaneous calcium events of VIP interneurons between pre- and post-drug period. The change was calculated as the post-drug event rate minus pre-drug event rate, divided by the pre-drug event rate. n = 53 cells from 6 animals. **(H)** Similar to (G) for SST interneurons. n = 117 cells from 7 animals. **(I)** Similar to (G) for PV interneurons. n = 135 cells from 5 animals. ** *P* < 0.01. Mixed effects model.

To evaluate the two mechanisms, we first measured the drug-evoked activity change in VIP interneurons. We initially tried using the Neuropixels and opto-tagging approach, but were unable to reliably find opto-tagged units in *Vip^Cre^*;Ai32 mice (n = 7 animals, 616 cells with no opto-tagged unit; data not shown). We therefore pursued an alternative strategy using two-photon microscopy to image somatic calcium transients. To express a genetically encoded calcium indicator in VIP interneurons, we injected AAV-CAG-Flex-GCaMP6f into the medial frontal cortex of *Vip^Cre^* mice (*33*). Through a chronically implanted glass window, we imaged GCaMP6f-expressing VIP interneurons in layer 2/3 of the medial frontal cortex of awake, head-fixed mice (**Figure 2B**). We imaged for 30 minutes, administered psilocybin (1 mg/kg, i.p.) or saline, and then imaged for another 90 minutes. Each mouse would receive both psilocybin and saline, delivered one week apart with the order randomized in a counter-balanced design. From the time-lapse images, we extracted fluorescence transients from each cell body and used an automated algorithm to quantify the calcium event rate for each cell (see **Methods**). **Figure 2C** shows the somatic calcium transients from three VIP interneurons before and after the administration of saline or psilocybin.

We imaged 6 *Vip^Cre^* animals (2 males, 4 females) to measure from 53 VIP interneurons (range: 4–11 cells per animal). We did not detect any effect of psilocybin on the calcium event rates in VIP interneurons (psilocybin: −3.3*±*2.2%; saline: −2.8*±*2.6%; main effect of treatment: *P* = 0.8, mixed effects model; **Figure 2D and 2G**). For completeness, we also imaged somatic calcium transients from SST and PV interneurons in layer 2/3 of the medial frontal cortex using *Sst^Cre^*and *Pvalb^Cre^* mice, respectively (**Figure 2E and 2F**). In line with our electrophysiology results, we observed a reduced rate of calcium events in SST interneurons following psilocybin administration (psilocybin: −10.5*±*6.7%; saline: 26.1*±*13.3%; n = 117 cells from 7 animals including 5 males, 2 females; main effect of treatment: *P* = 0.003, mixed effects model; **Figure 2H**). However, we did not see appreciable changes in calcium event rate in PV interneurons following psilocybin administration (psilocybin: 15.8*±*8.8%; saline: 15.0*±*8.1%; n = 135 cells from 5 animals including 3 males, 2 females; main effect of treatment: *P* = 0.98, mixed effects model; **Figure 2I**). The discrepancy between the electrophysiological and imaging results for PV interneurons is likely because two-photon imaging recorded from layer 2/3 cells, missing the deeper neurons that were preferentially captured by the Neuropixels probe (**Figure S4**). Nevertheless, the imaging data demonstrate that VIP interneurons did not have appreciable change in activity after psilocybin administration. Therefore, VIP interneurons are unlikely the reason for the psilocybin-induced suppression of SST interneurons.

### 5-HT_1A_ receptors gate psilocybin’s suppressive effect on SST interneurons

SST interneurons express 5-HT_1A_ receptors, which are G_i_-coupled receptors that should reduce neuronal excitability (*40, 41*). To determine if the 5-HT_1A_ receptor is involved in psilocybin’s drug action, we used the same Neuropixels and opto-tagging approach to record from SST interneurons in the medial frontal cortex of *Sst^Cre^*;Ai32 mice, with an additional step to inject the 5-HT_1A_ receptor antagonist WAY-100635 (2 mg/kg, i.p., administered via a catheter) 10 minutes prior to psilocybin administration (1 mg/kg, i.p., administered via a second catheter) or saline (**Figure 3A and 3B**). For saline, we recorded from 6 animals (4 males, 2 females) to obtain 1582 cells including 22 opto-tagged SST interneurons (range: 107-599 cells and 1-8 opto-tagged units per animal). The mean firing rates for frontal cortical SST interneurons were 6.9±1.0 Hz and 5.9±1.0 Hz before and after saline, respectively (**Figure 3C**). For psilocybin, we recorded from 6 animals (5 males, 1 female) to obtain 692 cells including 33 opto-tagged SST interneurons (range: 58–145 cells and 1–8 opto-tagged units per animal). In these animals pretreated with the 5-HT_1A_ receptor antagonist, psilocybin no longer reduced the firing of cortical SST interneurons (pre-psilocybin: 7.4±1.2 Hz, post-psilocybin: 7.9±1.0 Hz; **Figure 3D**). In fact, there was now a slight but significant increase in spike rate of SST interneurons after WAY-100635 and psilocybin (7.8±2.1%, mean ± SEM; main effect of treatment: *P* = 0.028, mixed effects model). The results support a role for the 5-HT_1A_ receptor. However, the experiment has a couple caveats. Although WAY-100635 is a well-characterized 5-HT_1A_ receptor antagonist (*42*), the compound has off-target effect at the dopamine D4 receptor (*43*). Moreover, WAY-100635 was administered systemically, so it can affect numerous cell types including cortical excitatory neurons that also express 5-HT_1A_ receptors.

**Figure 3.**
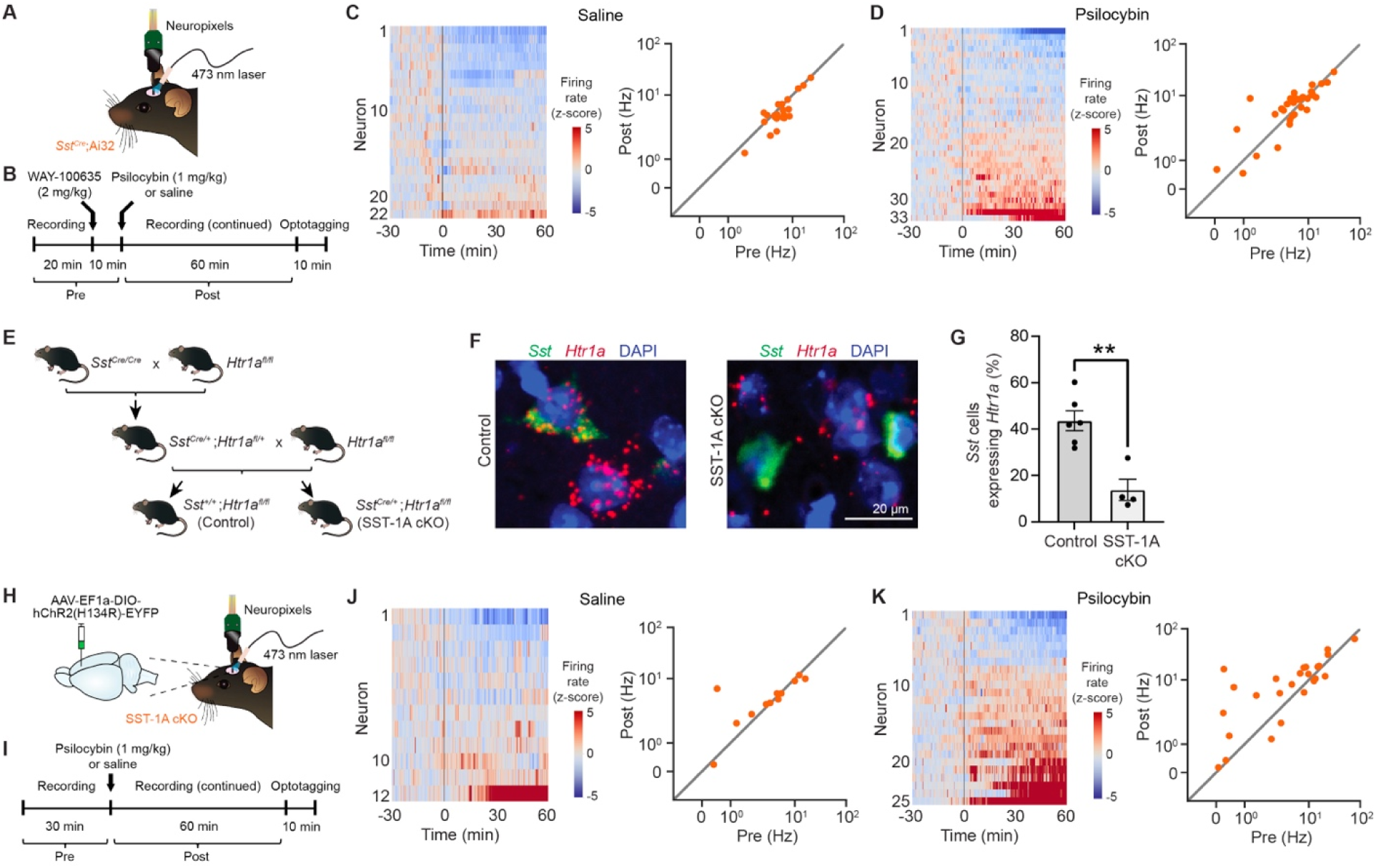
5-HT_1A_ receptors are responsible for psilocybin’s suppressive effects on SST interneurons. **(A)** Spiking activity of ChR2-expressing SST interneurons in *Sst^Cre^*;Ai32 mice was recorded using a Neuropixels probe. **(B)** The experimental timeline includes pre-drug (−30 – 0 min) and post-drug (0 – 60 min) periods. The 5-HT_1A_ receptor antagonist WAY-100635 was administered 10 minutes prior to psilocybin or saline injection. **(C)** Left, the activity of opto-tagged SST interneurons before and after saline administration in animals pretreated with WAY-100635. The firing rate of each neuron was converted to a z-score by normalization based on its pre-drug firing rate. Right, mean firing rates during pre- and post-drug periods for the opto-tagged SST interneurons. The axes use a symmetric logarithmic scale, which includes a linear region (0 to 0.03 Hz) that transitions to a logarithmic region (>0.03 Hz). Each dot represents one neuron. n = 22 cells from 6 animals. **(D)** Similar to (C) for psilocybin administration. n = 33 cells from 6 animals. **(E)** Breeding scheme to produce SST-1A cKO mice (conditional knockout of *Htr1a* in SST interneurons) and littermate controls. **(F)** Example images of *Sst* (green) and *Htr1a* (red) transcripts via RNAscope fluorescence in situ hybridization along with DAPI stain (blue) in control and SST-1A cKO mice. **(G)** Quantification of the RNAscope validation: fraction of *Sst*-expressing cells that contain *Htr1a* transcripts in the frontal cortex of control and SST-1A cKO mice. Each dot represents an individual animal. n = 1626 SST interneurons from 6 control animals, and 1396 SST interneurons from 4 SST-1A cKO mice. Two-tailed Wilcoxon rank-sum test. **(H)** AAV was used to induce expression of ChR2 in SST interneurons in the medial frontal cortex of SST-1A cKO mice. Spiking activity of ChR2-expressing SST interneurons was recorded using a Neuropixels probe. **(I)** The experimental timeline includes pre-drug (−30 – 0 min) and post-drug (0 – 60 min) periods. **(J)** Left, the activity of opto-tagged SST interneurons before and after saline administration in SST-1A cKO mice. The firing rate of each neuron was converted to a z-score by normalization based on its pre-drug firing rate. Right, mean firing rates during pre- and post-drug periods for the opto-tagged SST interneurons. The axes use a symmetric logarithmic scale, which includes a linear region (0 to 0.03 Hz) that transitions to a logarithmic region (>0.03 Hz). Each dot represents one neuron. n = 12 cells from 6 animals. **(K)** Similar to (J) for psilocybin administration. n = 25 cells from 11 animals. ** p < 0.01.

To test further the role of the 5-HT_1A_ receptor, we leveraged a *Htr1a^fl/fl^* mouse line that was generated in a prior study (*44*). We generated *Sst^Cre/+^*;*Htr1a^fl/fl^*mice (henceforth referred to as SST-1A cKO) and *Sst^+/+^*;*Htr1a^fl/fl^* littermates as controls (**Figure 3E**). The original study validated the conditional knockout using immunohistochemistry and slice electrophysiology (*44*). To characterize the knockout in our experimental context, we used RNAscope fluorescence in situ hybridization to visualize *Htr1a* and *Sst* transcripts around DAPI-positive nuclei in the medial frontal cortex (**Figure 3F**). In control animals, 44±4% of *Sst*-expressing cells contained *Htr1a* transcripts (mean ± SEM; n = 1626 SST cells from 6 animals including 4 females, 2 males; **Figure 3G**). This percentage is close to the value derived from the single-cell sequencing data, which showed *Htr1a* transcript in 39% of the frontal cortical SST interneurons (**Figure 2A**). By contrast, in SST-1A cKO animals, only 14±5% of *Sst*-expressing cells contained *Htr1a* transcripts (n = 1396 SST cells from 4 animals including 3 males, 1 female; *P* = 0.0095, two-tailed Wilcoxon rank-sum test). There are a couple reasons why the knockout was substantial but incomplete. Because the *Htr1a* gene lacks introns, the floxed construct placed loxP sites flanking the entire coding sequence, spanning a large distance of 1.3 kb. In addition, the chromatin state in frontal cortical SST interneurons may limit Cre recombinase access to the loxP sites. Both factors could contribute to reduced Cre-mediated recombination efficiency. The characterization confirmed that the *Htr1a^fl/fl^*conditional knockout mice can be used for cell type-specific manipulations in the mouse medial frontal cortex.

We injected AAV-EF1a-DIO-hChR2(H134R)-EYFP into the medial frontal cortex to express ChR2 selectively in SST interneurons of SST-1A cKO animals (**Figure 3H**). We used the Neuropixels and opto-tagging approach to record from SST interneurons, which now lacked *Htr1a* expression (**Figure 3I**). For saline, we recorded from 6 animals (2 males, 4 females) to obtain 886 cells including 12 opto-tagged SST interneurons (range: 76-261 cells and 1-4 opto-tagged units per animal). There was no effect of saline on the mean firing rate of SST interneurons in SST-1A cKO mice (pre-saline: 5.2±1.4 Hz, post-saline: 5.1±1.0 Hz; **Figure 3J**). For psilocybin, we recorded from 11 animals (7 males, 4 females) to obtain 927 cells including 25 opto-tagged SST interneurons (range: 17–200 cells and 1–5 opto-tagged units per animal). In mice with cell type-specific deletion of 5-HT_1A_ receptors, psilocybin had no effect on the activity of SST interneurons in the medial frontal cortex (pre-psilocybin: 10.4±3.2 Hz, post-psilocybin: 13.2±2.8 Hz; main effect of treatment: *P* = 0.4, mixed effects model; **Figure 3K**). Thus, results from the pharmacological blockade and conditional knockout experiments converge to pinpoint the 5-HT_1A_ receptor on SST interneurons as an important target for psilocybin’s drug action on SST interneurons.

### Frontal cortical SST interneurons contribute to psilocybin’s long-term effect on stress-related behavior

We next asked if the 5-HT_1A_ receptors on SST interneurons may play a role in psilocybin’s acute and long-term behavioral effects. We tested SST-1A cKO and control mice on three behavioral assays, with the tests spaced at least 2 weeks apart to allow for drug washout. Fear extinction is a preclinical behavioral assay that provides insight into stress-related psychopathology (*45, 46*). Psilocybin reduces freezing not only in the extinction session that immediately follows, but also in the extinction retention and fear renewal sessions days later (*47*). For the current experiment, on day 1, we paired auditory tones with foot shocks for fear conditioning in context A (**Figure 4A**; see **Methods**). On day 3, we administered psilocybin (1 mg/kg, i.p.) or saline, and then 30 minutes later had the first extinction session where only auditory tones were presented in context B. On day 4, we tested extinction retention by performing a second extinction session in context B. Finally, on day 12, we tested fear renewal, which is an extinction session performed in context C. Both SST-1A cKO and control mice readily acquired conditioned freezing (main effect of tones: *P* < 0.001, mixed-effects model; n = 14–20 mice per group; **Figure 4B**). In control mice, a single dose of psilocybin (1 mg/kg, i.p.) significantly reduced freezing during fear extinction compared to saline treatment. However, this psilocybin-facilitated extinction was absent in SST-1A cKO mice (interaction effect of treatment and genotype: *P* = 0.002, two-factor ANOVA; **Figure 4C**). Similarly, psilocybin’s effect on freezing during extinction retention was abolished in SST-1A cKO mice (interaction effect of treatment and genotype: *P* = 0.001, two-factor ANOVA; **Figure 4D**). There was no interaction effect detected for fear renewal (main effect of treatment: *P* = 0.004, main effect of genotype: *P* = 0.004, two-factor ANOVA; **Figure 4E**).

**Figure 4.**
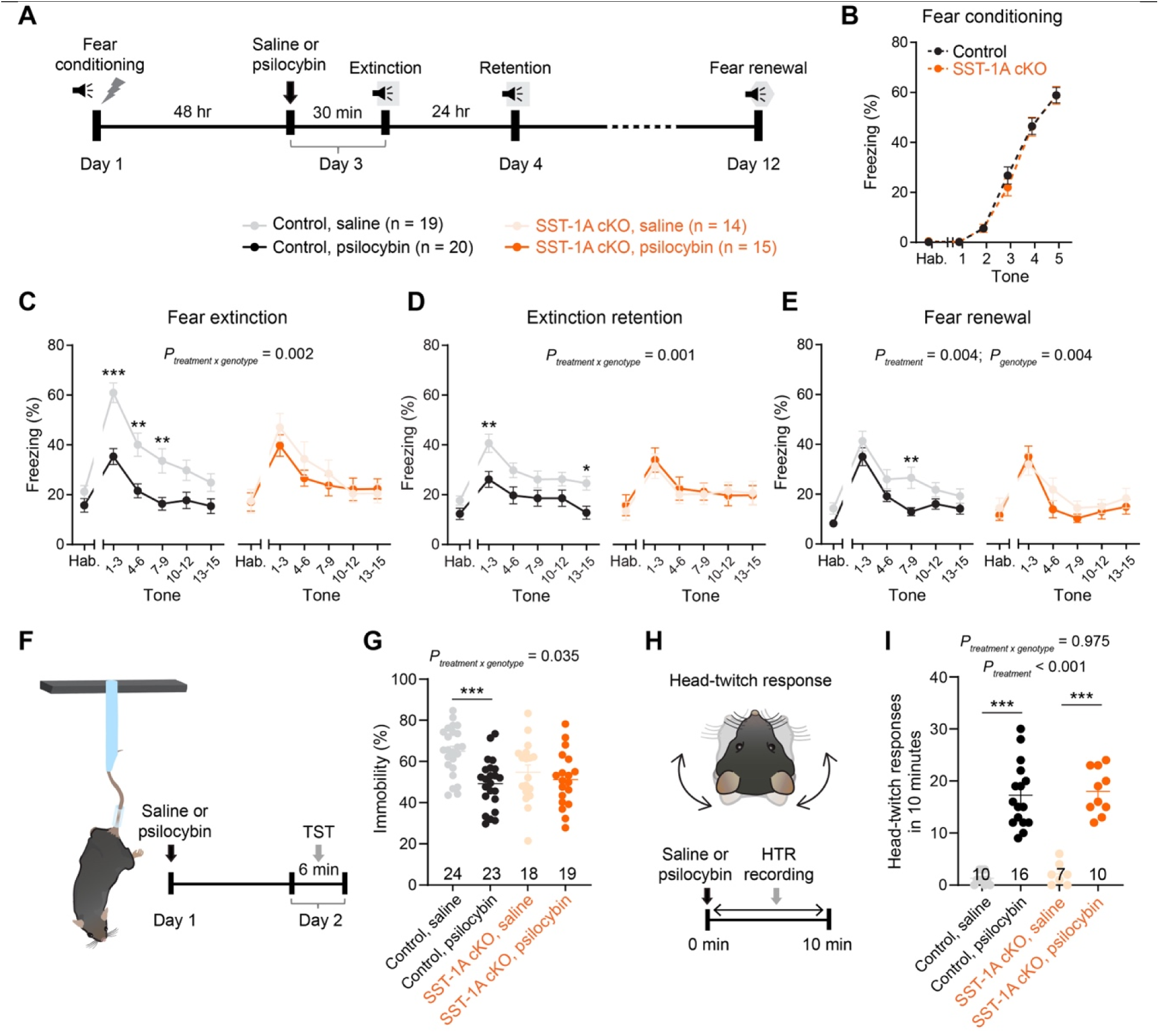
5-HT_1A_ receptor knockout in SST interneurons influences psilocybin’s effects on stress-related behaviors. **(A)** Experimental timeline of fear conditioning and extinction tests. **(B)** Percentage of time spent freezing during fear conditioning of control (black) and SST-1A cKO mice (orange). **(C)** Percentage of time spent freezing during fear extinction measured in control mice after saline (gray) or psilocybin (black) and in SST-1A cKO mice after saline (light orange) or psilocybin (dark orange). n = 19 mice (control, saline), 20 mice (control, psilocybin), 14 mice (SST-1A cKO, saline), 15 mice (SST-1A cKO, psilocybin). Two-factor ANOVA. **(D)** Similar to (C) for extinction retention. **(E)** Similar to (C) for fear renewal. **(F)** Schematic of the tail suspension test. **(G)** Percentage of time spent immobile in control mice after saline (gray) or psilocybin (black) and in SST-1A cKO mice after saline (light orange) or psilocybin (dark orange). Each dot represents an individual animal. n = 24 mice (control, saline), 23 mice (control, psilocybin), 18 mice (SST-1A cKO, saline), 19 mice (SST-1A cKO, psilocybin). Two-factor ANOVA. **(H)** Schematic of the head twitch response. **(I)** Head twitch response count within 10 min after psilocybin or saline administration in control mice after saline (gray) or psilocybin (black) and in SST-1A cKO mice after saline (light orange) or psilocybin (dark orange). Each dot represents an individual animal. n = 10 mice (control, saline), 16 mice (control, psilocybin), 7 mice (SST-1A cKO, saline), 10 mice (SST-1A cKO, psilocybin). Two-factor ANOVA. Data present means ± SEM across mice. * p < 0.05; ** p < 0.01; *** p < 0.001, *post hoc* with Bonferroni correction for multiple comparisons.

The tail suspension test measures stress-induced behavioral despair and is used for antidepressant screening (*48*) (**Figure 4F**). Psilocybin significantly reduced immobility time 24 hours after administration compared to saline treatment in control mice, but this effect was absent in SST-1A cKO animals (interaction effect of treatment and genotype: *P* = 0.035, two-factor ANOVA; n = 18–24 per group; **Figure 4G**). We note that saline-treated SST-1A cKO animals had a low level of immobility, which may occluded detection of any further decrease induced by psilocybin. This resilient phenotype at baseline for the SST-1A cKO animals is consistent with earlier work demonstrating that constitutively disinhibiting SST interneurons leads to an antidepressive-like state (*49*).

Finally, the head-twitch response is an assessment of the acute drug action of psychedelics, with the head-twitch potency of a compound in mice relating to its hallucinogenic potency in humans (*50*) (**Figure 4H**). As expected, psilocybin induced robust increase in the number of head twitches in control mice, and this acute behavioral effect was unaffected by the knockout of 5-HT_1A_ receptors in SST interneurons (interaction effect of treatment and genotype: *P* = 0.98, two-factor ANOVA; n = 7–16 per group; **Figure 4I**). Collectively, these results demonstrate that the 5-HT_1A_ receptors on SST interneurons modulate psilocybin’s long-term effects on stress-related behaviors, but do not contribute to its acute head-twitch response in mice.

## DISCUSSION

This study reveals that psilocybin differentially affects the firing activity of GABAergic cell populations. In the medial frontal cortex, psilocybin reduced the spiking activity of SST interneurons while increasing the firing of PV interneurons. Pharmacological blockade and conditional knockout experiments demonstrate that the psilocybin-induced suppression of SST interneurons is mediated by the 5-HT_1A_ receptors expressed on the SST interneurons. This modulation of GABAergic inhibition is functionally important, because deleting the 5-HT_1A_ receptors from SST interneurons impaired the long-term ameliorating effects of psilocybin on stress-related behaviors.

In adult animals, learning is accompanied by a dynamic reorganization of cortical inhibitory tone, marked by cell type-specific changes in interneuron activity (*51, 52*). In particular, SST interneurons target the dendrites of pyramidal cells to regulate excitability (*53*). Transient reduction of SST interneuron activity and the resulting disinhibition have emerged as an important mechanism gating synaptic plasticity during learning (*54, 55*), including for the encoding of fear memory (*56*). Our results suggest that psychedelics may leverage a similar dendritic disinhibition mechanism. We showed that psilocybin suppresses the activity of SST interneurons in this study. The expected downstream consequence is increased dendritic excitability, consistent with recent report of an elevated rate of dendritic calcium transients in the pyramidal tract subtype of pyramidal cells after psilocybin administration (*57*). Overall, our findings support a model in which psilocybin shifts inhibition along the somatodendritic axis of cortical pyramidal cells: suppressing SST interneurons to reduce dendritic inhibition, while activating PV interneurons to increase perisomatic inhibition and restrict excessive spiking output.

We focused the current study on the medial frontal cortex, because this mouse brain region responds robustly to psilocybin administration, as identified by brain-wide c-Fos mapping (*58*). However, the effects of psychedelics are likely to vary across brain regions. For example, psychedelics affect the physiology of the hippocampus, which contains microcircuits composed of PV, SST, and VIP interneurons that are analogous to those in the neocortex. PV interneurons in the ventral hippocampus express 5-HT_2A_ receptors, which are critical for mediating the acute anxiolytic effects of psychedelics (*59*). In contrast to the cortex, hippocampal SST interneurons appear to express also 5-HT_2A_ receptors preferentially (*60*), contributing to slow oscillatory activity (*61*). Even within cortical regions, psychedelic effects may vary among finer interneuron subtypes. SST interneurons can be subdivided into at least 8 subtypes based on genetic markers and morphological features (*62, 63*). Our results indicate that psilocybin does not uniformly suppress the activity of all SST interneurons. Moreover, when 5-HT_1A_ receptor was blocked pharmacologically, psilocybin increased the activity of SST neurons, possibly because a fraction of the interneurons expresses 5-HT_2A_ receptors (*64*) and their influence was unmasked by the manipulation. highlighting the need for future studies to classify cell type-specific responses to psychedelic drugs in greater detail.

We did not detect psilocybin-evoked activity changes in frontal cortical VIP interneurons. VIP interneurons express the G_q_-coupled 5-HT_2C_ receptors, but their effect on cellular excitability can vary by brain region and cell type, as shown in studies of the amygdala (*65*), claustrum (*66*), ventral hippocampus (*67*), and piriform cortex (*68*). Several possibilities could explain the lack of effect in frontal cortical VIP interneurons. The G_q_-coupled pathway in cortical VIP interneurons may not engage ion channels to alter excitability. In addition, VIP interneurons express abundant 5-HT_3_ receptors, which are normally activated by endogenous serotonin to increase excitability (*69*). Classic psychedelics silence the serotonergic neurons in the dorsal raphe (*70, 71*), potentially diminishing serotonergic drive onto VIP interneurons. This could mask any excitatory effects mediated through the 5-HT_2C_ receptors, resulting in an overall lack of activity change.

In summary, our findings illuminate how specific receptors and GABAergic cell types within the cortex orchestrate the complex effects of psilocybin. The results are consistent with psilocybin causing a coordinated shift in the inhibitory tone along the somatodendritic axis of pyramidal cells. Interestingly, NMDA receptor antagonists such as the rapid acting antidepressant ketamine also induce dendritic disinhibition (*72*). However, NMDAR receptor antagonists additionally reduce perisomatic inhibition (*73*), which differs sharply from the increased PV interneuron activity after psilocybin administration. Future studies will continue to uncover these distinct and shared microcircuit-level mechanisms engaged by different classes of drugs. By dissecting the cell type-specific responses to psilocybin, the present study lays the groundwork for understanding how different cellular components work in concert to drive the acute and long-term effects of psychedelics.

## SUPPLEMENTARY MATERIALS

Supplementary materials include Figure S1 – S5.

## ACKNOWLEDGEMENTS

We thank Eirene Markenscoff-Papadimitriou for help with RNAscope and Olesia Bilash for comments on the manuscript. Psilocybin was provided by Usona Institute’s Investigational Drug & Material Supply Program; the Usona Institute IDMSP is supported by Alexander Sherwood, Robert Kargbo, and Kristi Kaylo in Madison, WI. The authors acknowledge the support of NIH/NIMH grants R01MH121848, R01MH128217, R01MH137047 (A.C.K.), One Mind–COMPASS Rising Star Award (A.C.K.), NIH training grants T32GM007205 (P.A.D. and N.K.S.), NIH fellowships F30DA059437 (P.A.D.), and F30MH129085 (N.K.S.), and NIH instrumentation grant S10OD032251 (Cornell Biotechnology Resource Center Imaging Facility).

## AUTHOR CONTRIBUTIONS

P.A.D. and A.C.K planned the study. P.A.D., Q.J., and C.A.K. conducted the electrophysiological experiments. P.A.D. analyzed the electrophysiological data. N.K.S. and J.W. performed the imaging experiments and analyzed the data. L.S. and C.L. performed the behavioral experiments and analyzed the data. A.M.W. performed the RNAscope experiment. Q.J., C.W.C., and J.D.N. assisted in histology. T.S. generated and provided the *Htr1a^f/f^* mice. P.A.D. and A.C.K. drafted the manuscript. All authors reviewed the manuscript before submission.

## METHODS

### Animals

Experimental procedures were approved by the Institutional Animal Care & Use Committees at Cornell University and Yale University. C57BL/6J (#000664), *Sst^Cre^*(#028864), *Pvalb^Cre^* (#017320), *Vip^Cre^* (#031628), and Ai32 (#024109, all from Jackson Laboratory) mice were bred in our animal facility. The *Htr1a^fl/fl^* strain was generated for a prior study (*44*), then transferred and bred in our animal facility. To generate mice with *Htr1a* deletion in SST interneurons, *Sst^Cre/Cre^* mice were bred with *Htr1a^fl/fl^*mice. This cross produced *Sst^Cre/+^*;*Htr1a^fl/+^*progeny that was bred with *Htr1a^fl/lf^* mice to generate the *Sst^Cre/+^*;*Htr1a^fl/fl^* (SST-1A cKO) mice and *Sst^+/+^*;*Htr1a^fl/fl^* littermate control mice used for experiments. For electrophysiology, 8 to 10-week-old *Sst^Cre^*;Ai32, *Pvalb^Cre^*;Ai32, *Vip^Cre^*;Ai32, and SST-1A cKO mice were used. For two-photon calcium imaging, 8 to 10-week-old *Sst^Cre^*, *Pvalb^Cre^*, and *Vip^Cre^* mice were used. Heterozygous mice were used to minimize potential effects of hypomorphic allele in homozygous mice. Animals were housed in groups with 2 – 5 mice per cage in a temperature-controlled room, operating in normal 12 hr light - 12 hr dark cycle (8:00 AM to 8:00 PM for light). Food and water were available ad libitum. Animals were randomly assigned to different experimental groups.

### Drugs and viruses

Non-GMP psilocybin was obtained through the Investigational Drug & Material Supply Program at Usona Institute. The chemical composition was confirmed by HPLC at Usona Institute. WAY-100635 (#4380, Tocris Biosciences) was administered at a dose of 2 mg/kg. AAV1-CAG-Flex-GCaMP6f-WPRE-SV40 (Catalog #100835) and AAV1-EF1a-DIO-hChR2(H134R)-EYFP-WPRE-HGHpA (Catalog #20298) were purchased from Addgene. All viruses had titers ≥ 7×10^12^ vg/mL. The viruses were stored at −80°C. Before stereotaxic injection, they were taken out of the −80°C freezer, thawed on ice, and diluted to the corresponding titer for injection.

### Surgery

Prior to surgery, each mouse received injections of dexamethasone (3□mg/kg, i.m.; #1DEX022, Bimeda) and carprofen (5□mg/kg, s.c.; #059149, Covetrus) for anti-inflammatory and analgesic purposes. Anesthesia was induced with 2–3% isoflurane and the mouse was positioned in a stereotaxic apparatus (Model 900, David Kopf Instruments). Anesthesia was maintained with 1–1.5% isoflurane throughout the procedure. Body temperature was kept at 38°C using a far-infrared warming pad (#RT-0515, Kent Scientific). Petrolatum ophthalmic ointment (Optixcare Eye Lube, #062143, Covetrus North America) was applied to protect the eyes. The hair on the head was shaved. An incision was made to remove the skin and the periosteum was cleared. The scalp was disinfected by wiping with ethanol pads and povidone-iodine. We targeted the recording location to be ACAd and medial MOs subregion of medial frontal cortex (AP: +1.7□mm, ML: −0.25□mm, relative to bregma). To reach the target, we would center the craniotomy at ML = −0.5 mm, with intent to insert the probe in the medial portion of the craniotomy. We would mark the spot for the center of craniotomy with ink (e.g., skin-safe pen). A small burr hole was made above the cerebellum using a 0.7 mm burr (#19007-07, Fine Science Tools) and a handheld dental drill (#HP4-917, Foredom). A 0.86 mm self-tapping bone screw (#19010-10, Fine Science Tools) was placed through the skull bone into the cerebellum to serve both as a ground and to provide mechanical support for the head plate implant. A custom stainless steel head plate (fabricated by eMachineShop; design available at https://github.com/Kwan-Lab/behavioural-rigs) was affixed onto the skull using a quick adhesive cement system (Metabond, #S396, Parkell) while leaving a 5-mm diameter area clear around the planned recording site and also leaving sufficient metal contact in the screw to make contact with reference wire in the future. Any area not covered be Metabond was then covered with silicone elastomer (#10006546, Smooth-On, Inc.). At the end of surgery, animal was given carprofen (5□mg/kg, s.c.) immediately and then again once on each of the following 3 days. The mouse would recover for at least 1 week after surgery before the recording day.

Specifically for electrophysiology with *SST^Cre/+^*;*Htr1a^fl/fl^*mice, we required virally mediated expression of ChR2 in SST interneurons. An additional surgery was performed. During this surgery, instead of marking or denting the spot for the medial frontal cortex, a burr hole was made above the ACAd and medial MOs subregion of medial frontal cortex using the burr and handheld dental drill. AAV1-EF1a-DIO-hChR2(H134R)-EYFP-WPRE-HGHpA virus was delivered intracranially into the brain by inserting a pipette pulled from a borosilicate glass capillary and using an injector (Nanoject II Auto-Nanoliter Injector, Drummond Scientific).

Injections were done using 4.6□nL pulses with 20□s interval between each pulse. To reduce backflow of the virus, we waited 5–10 min after completing an injection at one site before retracting the pipette to move on to the next site. A total of 4 injections (totaling 200 nL) sites were targeted, corresponding to the vertices of a 0.2 mm-wide square centered at the coordinates for the targeted brain region. Throughout the procedure, the brain surface was kept moist with artificial cerebrospinal fluid (ACSF; in mM: 135 NaCl, 5 HEPES, 5 KCl, 1.8 CaCl2, 1 MgCl2; pH: 7.3). After injections, the craniotomies were covered with silicone elastomer. The skin was sutured (#1265B, Surgical Specialties Corporation). Mice would recover for at least 7 day prior to the surgery to implant the head plate, and we would wait a minimum of 3 weeks to allow for viral-mediated expression.

For two-photon imaging, surgeries were performed involving the same pre- and post-operative procedures. A burr hole was made to inject 92 nL of AAV1-CAG-Flex-GCaMP6f-WPRE-SV40 (1:20 diluted in PBS) into the ACAd and medial MOs subregion of medial frontal cortex (AP: 1.5□mm, ML: −0.4□mm, DV: −1.0□mm). After 2–3 weeks, the mouse underwent a second procedure, with the same pre- and post-operative procedures, to implant a glass window for imaging. An incision was made to expose the skull, and the surface was cleaned to remove connective tissues. The dental drill was used to make a ∼3-mm-diameter circular craniotomy above the previously targeted location at the medial frontal cortex. ACSF was used to immerse the exposed dura in the craniotomy. A two-layer glass window was assembled by bonding two round coverslips (3 mm diameter, 0.15 mm thickness; #640720, Warner Instruments) via ultraviolet light-curing optical adhesive (#NOA 61, Norland Products) using an ultraviolet illuminator (#2182210, Loctite). The glass window was placed over the craniotomy and, while maintaining a slight pressure, super glue adhesive (Henkel Loctite 454) was carefully used to secure the window to the surrounding skull. A stainless steel headplate (eMachineShop; design available at https://github.com/Kwan-Lab/behavioral-rigs) was secured on the skull and centered on the glass window using a quick adhesive cement system (Metabond, Parkell). Mouse would recover for at least 10 days prior to imaging.

### Electrophysiology

The electrophysiology rig was built on an air table (63-9012E, TMC) inside a Faraday cage (81-333-06, TMC). Mice were habituated to head fixation with increasing duration over several days. At least 3 hr before recording, mice were anesthetized with isoflurane. The silicone elastomer covering was removed. The exposed skull was sterilized with alternating wipes of 70% ethanol and betadine. A 2-mm-diameter craniotomy was made at the previously marked spot of the medial frontal cortex using the dental drill equipped with the 0.7-mm-diameter burr. Cold (4°C) ACSF was applied intermittently throughput the procedure to irrigate the area, clear debris, and reduce heat generated by drilling. Care was taken to minimize bleeding and keep the area clear of bone fragments. Using a tungsten needle (#10130-10, Fine Science Tools) held by a pin holder (#26018-17, Fine Science Tools), we would make a small slit in the dura to facilitate the insertion of the Neuropixels probe. To protect the exposed brain tissue and maintain moisture, a piece of ACSF-soaked Surgifoam (#1972, Johnson & Johnson Ethicon) was placed over the surface and sealed with silicon polymer (#0318, Smooth-On, Inc.). A 22-gauge intravenous catheter system (#B383323, BD Saf-T-Intima Closed IV Catheter Systems) was prepared by first removing the plastic wings and pre-filled with psilocybin or saline and maintained at a neutral pressure. We loaded the drug into the catheter first before inserting the catheter to avoid air bubbles in the system. For WAY-100635 experiments, the antagonist solution was preloaded into a second catheter, so eventually the mouse would have two catheters inserted. At 1 hr prior to recording, mice were briefly anesthetized with isoflurane via a nose cone, positioned supine, and implanted with the intravenous catheter to their intraperitoneal cavity. The catheter was inserted at a shallow angle at ∼5 mm lateral from the midline at the level of hip joint into the intraperitoneal cavity. The base of the needle was fixed with a few drops of Vetbond tissue adhesive (#1469SB, 3M). The adhesive would dry completely and, once dried, the needle was carefully withdrawn by pulling back the cylindrical portion at the end of the catheter was stabilizing the base. The mice were then head fixed and the catheter tubing and the tail of the mouse were secured with tape. The silicon polymer and Surgifoam were removed from the craniotomy and the exposed brain tissue was filled with ACSF. A high-density Neuropixels 1.0 probe (#PRB_1_4_0480_1_C, IMEC) with the external ground and reference shorted with a silver wire. The ground wire was twisted around the ground screw, ensuring there was a good contact. The probe was coated using a 10 μL drop of CM-DiI (1 mM in ethanol, prepared by putting 50 μg of CM-DiI into 50 μL of 100% ethanol; #C7000, Invitrogen). This step involves first dipping a pipette into isopropyl alcohol (AB07015, AmericanBio), drawing up 10 μL of the solution and then ejecting to create a bubble, which was moved up and down along the probe to “paint” the probe until about half of the bubble has evaporated, taking care to not touch the probe with the pipette. The probe was positioned to the surface of the brain above the craniotomy using a micromanipulator on an inverted arm (MPM-4 DOF Arm-Inverted, New Scale Technologies), mounted in a ring platform system (M3-LS-3.4-15-XYZ-MPM-Inverted, New Scale Technologies), controlled via a joystick (Extreme 3D Pro, Logitech) under visual guidance using a digital microscope with extra-long working distance and magnification (AM73115MTF, Dino-Lite Edge 3.0). The z-position was zeroed at the point of first contact between the probe and the brain surface. The probe was then slowly lowered at a speed of 1.67 μm/s into the brain, to the target depth of 2000 μm below the surface of the brain. At the target depth, we waited for the probe to settle for at least 30 min before recording began. The probe was configured to record from the bottom 384 sites closest to the probe tip, which was the default configuration. The sites for Neuropixels 1.0 probe were arranged in 2 columns and spaced 20 μm apart, so the sites spanned 3840 μm. Our targeted depth was 2000 μm, resulting in only the lower subset of recording sites positioned within the brain. The more dorsally positioned recording sites were active but located outside the brain and therefore did not record neural activity. Data were acquired using a chassis system (PXIe-1071, National Instruments) containing data acquisition board (PXIe-8381, National Instruments) and the IMEC Neuropixels PXIe acquisition module card. The OpenEphys software was set to external reference mode. Action potentials were recorded at 30 kHz with gain of 500 and local field potentials were recorded at 2.5 kHz with gain of 250. Following the start of recording, 30 min of baseline activity was collected. The animal then received either psilocybin or saline via the catheter and recording continued for an additional 60 min. In a subset of experiments, the animal was additionally administered with WAY-100635 via a separate catheter 10 min prior to psilocybin administration. At the end of each recording session, optotagging was performed to identify ChR2-expressing neurons. A fiber-coupled 473 nm, 50 mW OBIS LX laser (#1193829, Coherent) was connected to a patch cable with a 200 μm core, 0.5 NA optical fiber (M104L01, ThorLabs), with an unjacketed end that was mounted on another micromanipulator on an inverted arm in the ring platform system. The optical fiber was moved to position above the medial frontal cortex aimed at the craniotomy. The OpenEphys software was used to trigger a PulsePal (#1102, Sanworks) to drive the analog input in the back panel of the OBIS laser control unit to produce pulses at 1 Hz and ∼25 mW/mm^2^ per trial. Typical pulse duration was 20 ms, but in a small subset of trials the duration was up to 50 ms. Each trial lasted for 1 s, with inter-trial interval of 950 - 980 ms, and at least 500 trials were conducted per session. This trigger from the OpenEphys was split and also routed to the trig input of the IMEC Neuropixels PXIe acquisition module card, so the timing of the laser stimulation can be aligned to the spike times.

After completion of an electrophysiological recording, mice were immediately perfused with PBS, followed by paraformaldehyde solution (PFA, 4% (v/v) in PBS). The brains were extracted and further fixed in 4% PFA at 4°C for 12 – 24 hr. Subsequently, 40-µm-thick coronal sections were obtained using a vibratome (#VT1000S, Leica) and mounted on slides including Vectashield containing DAPI (#H-1200-10, Vector Laboratories) with glass coverslips. Sections were imaged using a wide-field fluorescence microscope (BZ-X810, Keyence). The Neuropixels probe was cleaned between recordings by soaking in 1% Tergazyme solution (Alconox) overnight, then in DI water, and then isopropyl alcohol (AB07015, AmericanBio) before being allowed to air-dry prior to the next session.

### Analysis of electrophysiology data

We used the wrapper software SpikeInterface (*74*) to execute an analysis pipeline for preprocessing, spike sorting, and quality metric calculation. Preprocessing included (1) passing the data through a high pass filter with cutoff at 400 Hz, (2) detecting and removing bad sites based on noise level, which discard mostly the dorsally positioned recording sites that were not in the brain, (3) phase shift correction, to account for small delays between sites, and (4) common median reference subtraction, to reduce noise particularly the laser-induced artifacts, which were applied to the data from the entire session. Spikes were sorted and putative single units were identified along with drift correction via automated procedures using Kilosort 2.5 (*75*). The sorted spikes were then manually curated using Phy (https://github.com/kwikteam/phy). Quality metrics and waveform features were generated via SpikeInterface. The curated units were screened for four quality metrics: a presence ratio of ≥0.9, inter-spike interval violations ratio of <0.5, amplitude cutoff of <0.1, and isolation distance >20. Presence ratio measures the fraction of time during the recording in which a unit has at least one action potential, with time bin width of 60 s. Inter-spike-interval violations ratio measures the fraction of spikes of the unit that occurred in rapid succession within 1.5 ms, relative to the true spikes of the unit. Amplitude cutoff estimates the fraction of missed spikes, i.e. spikes of a unit that was below the spike detection threshold. Isolation distance calculates the distance between a unit and other units based on waveforms. Only manually curated units that additionally met all quality metrics were included for further analysis. To identify opto-tagged neurons, we created peri-stimulus time histograms by aligning spiking activity to the onset of laser stimulation. We classified opto-tagged neurons by considering the latency to spike and reliability of spiking in response to onset of laser stimulation. To compare firing rates during pre- versus post-drug periods, we generated scatter plots with the axes plotted in symlog scale, consisting of a linear region and a logarithmic region with the transition set at 0.03 Hz, using the ‘symlog’ function in Matplotlib with the linthresh parameter set to 0.03.

### Histology of electrophysiology data

After electrophysiological experiments, we performed histology and successfully reconstructed probe trajectory in most animals. Mice were perfused with PBS, followed by paraformaldehyde solution (PFA, 4% (v/v) in PBS). The brains were extracted and further fixed in 4% PFA at 4□°C for 12–24□h. Subsequently, 40-µm-thick coronal sections were obtained using a vibratome (VT1000S, Leica) and mounted onto slides with glass coverslips using Vectashield containing DAPI (H-1200-10, Vector Laboratories). The sections were imaged using a wide-field fluorescence microscope (BZ-X810, Keyence). To locate the Neuropixels probe, we used SHARP-TRACK (*76*) to align the fluorescence images of the coronal sections, including the DiI signals from the probe, with the mouse brain reference atlas in the Allen Common Coordinate Framework (*77*). To estimate the location of each unit, we used the channel with the largest voltage deflection in spike waveform. We located 12/12 and 27/27 opto-tagged cells for saline and psilocybin conditions for *Sst^Cre^*;Ai32 animals, and 22/26 and 27/27 opto-tagged cells for saline and psilocybin conditions for *Pvalb^Cre^*;Ai32 animals.

### Two-photon imaging

Two-photon imaging experiments were performed using a Movable Objective Microscope (MOM, Sutter Instrument) equipped with a resonant-galvo scanner (Rapid Multi Region Scanner, Vidrio Technologies) and a water-immersion 20X objective (XLUMPLFLN, 20x/0.95 N.A., Olympus). ScanImage 2020 software was used to control the microscope for image acquisition. To visualize GCaMP6f, we used an excitation wavelength of 920 nm and emission was collected behind a 475 – 550 nm bandpass filter. The laser power measured at the objective was typically ≤120 mW and varied depending on the imaging depth. When imaging the same field of view across days, the laser power was kept the same in each imaging session. Prior to imaging, the mouse was habituated to head fixation in an acrylic tube under the microscope for 3–4 days, increasing the duration each day. We targeted fields of view in the medial portion of the cranial window in order to record from neurons in ACAd and medial MOs. To examine the acute treatment effects, we imaged a single field of view for 30 min to obtain pre-treatment baseline data. Imaging was then paused to inject psilocybin (1 mg/kg, i.p.) or saline (10 mL/kg, i.p.). Almost immediately following injection (<1 min), we imaged the same field of view again for 90 min to acquire post-treatment data. Each animal received both psilocybin and saline, with at least 1 week between imaging sessions and the order of treatment balanced across subjects. The same field of view was imaged for each treatment, providing a precise within-cell estimate of the drug’s effect relative to saline. Only cells with robust calcium transients in both imaging sessions were kept for imaging analysis.

### Analysis of imaging data

The multi-page .tiff image files from each experiment were concatenated and processed with NoRMCorre in MATLAB to correct for non-rigid translational motion. Regions of interest (ROI) corresponding to cell bodies were manually traced using an in-house graphical user interface in MATLAB. For each ROI, the pixel-wise average was calculated at each data frame to generate a fluorescence time course F_ROI_(t). Since calcium imaging was performed on the same field of view before and after drug injection, a single ROI mask was used to extract calcium signals for all images in an experiment. Next, each ROI was processed to reduce the contribution from background neuropil. Taking each ROI’s area and considering a circle with equivalent area that has radius *r*, a neuropil mask specific to that ROI was created as an annulus with inner radius 2*r* and outer radius 3*r* centered on the centroid of the ROI. To exclude neuropil mask pixels that may belong to unselected cell bodies, we calculated the time-averaged signal for each pixel, taking the median amongst pixels in the mask. Pixels were excluded from the neuropil mask if their time-averaged signal was higher than the median. Finally, the remaining pixels in the neuropil mask were averaged per data frame to generate F_neuropil_(t). Each ROI had the fluorescence from its neuropil mask subtracted as follows:

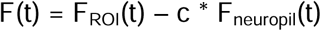

where the neuropil correction factor, c, was set to 0.4. Next, the fractional change in fluorescence ΔF/F(t) was calculated for each ROI by normalizing F(t) against its baseline, F_0_(t), estimated as the 10th percentile within a two-minute sliding window:

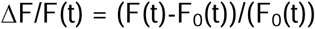

Calcium events were detected using automated procedure for each cell using a deconvolution “peeling” algorithm (*78*). The peeling algorithm uses an iterative template-matching procedure to decompose a ΔF/F(t) trace into a series of elementary calcium events. The template for elementary calcium events was defined on a cell-by-cell basis by calculating the median amplitude and single-exponential decay time constant from calcium transients exceeding 2 standard deviations of the cell’s mean signal. Briefly, the algorithm searches a given ΔF/F(t) trace for a match to the template calcium event, subtracts it from the trace (i.e., “peeling”), and successively repeats the matching process until no more event is found. This event detection process outputs the recorded event times with a temporal resolution by the original imaging frame rate. In this way, it is possible to detect multiple calcium events during the same imaging frame (e.g., for large-amplitude transients). For each imaging session, an ROI’s calcium event rate was computed by dividing the number of calcium events by the duration of the imaging session. The change in calcium event rate due to drug administration was computed for each ROI using the post-drug minus pre-drug values divided by the pre-drug values.

### RNAscope fluorescence *in situ* hybridization (FISH)

To quantify the efficiency of the knockout in the SST-1A cKO mice, RNAscope (Advanced Cell Diagnostics, ACD) was conducted using the Multiplex Fluorescent Reagent Kit v2 (#323270) and RNAscope probes for *Htr1a* (#312301-C2) and *Sst* (#404631). Mice were anesthetized with isoflurane and transcardially perfused with chilled phosphate buffered saline (PBS), followed by 4% paraformaldehyde (PFA) in PBS. Brains were extracted and fixed in 4% PFA for 24 h at 4*°*C then moved through a gradient of 10%, 20%, and 30% sterile-filtered sucrose at 4*°*C, allowing enough time to sink in each solution (16–20 h). Tissue was frozen in optimal cutting temperature (OCT) embedding medium (#4583, Tissue-Tek) and 14 μm coronal sections of the PFC were cut with a cryostat (Thermo Scientific Microm HM 550) and mounted onto ColorFrost Plus slides (#22-230-892, Fisher Scientific). Slides were stored at −80*°*C until the RNAscope pretreatment protocol and assay.

To pretreat the tissue, slides were washed in 1X PBS, baked on a slide warmer at 60*°*C for 45 min, and post-fixed in 4% PFA at 4*°*C for 15 min. Sections were dehydrated in 50%, 70%, and 100% ethanol for 5 min each at room temperature, then allowed to dry completely. Hydrogen peroxide (#322335, ACD) was applied to each section and allowed to sit for 10 min at room temperature. Slides were then rinsed in distilled water before incubating in 96*°*C Target Retrieval Buffer (#322000, ACD) for 8 min. Following a wash in distilled water and an additional dehydration in 100% ethanol for 3 min, slides were dried on the 60*°*C slide warmer and, once completely dry, barriers were drawn around each tissue section with an ImmEdge Hydrophobic Barrier Pen (#319918, ACD). After the barrier was completely dry, RNAscope Protease III (#322337, ACD) was added to each section and incubated at 40°C for 30 min in a HybEZ™ II Oven (#321711, ACD). Slides were rinsed in distilled water and immediately processed according to the instructions for the RNAscope assay.

The C2 (*Sst*) probe was diluted in the C1 (*Htr1a*) probe at a 1:50 ratio and 75-125 μl (depending on size of drawn hydrophobic barrier) of the mix was added to completely cover each section.

Sections were incubated with the probe mix at 40*°*C in the HybEZ™ II Oven for 2 h. After washing in 1X RNAscope Wash Buffer (#310091, ACD), slides were stored in 5X saline-sodium citrate (SSC) buffer overnight. On the second day of the protocol, TSA Vivid Fluorophores 520 (#323271, ACD) and 650 (#323273, ACD) were diluted at a 1:2000 ratio in TSA dilution buffer (#322810, ACD). The probe signal was amplified, and fluorophores were assigned to each channel using enough of each detection reagent (#323110, ACD) and diluted fluorophore to completely cover each tissue section. Solutions were applied for specific incubation times in the following order at 40□C in the HybEZ™ II Oven: AMP 1 (30 min), AMP 2 (30 min), AMP 3 (15 min), HRP-C1 (15 min), TSA Vivid Fluorophore 650 (30 min), HRP blocker (15 min), HRP-C2 (15 min), TSA Vivid Fluorophore 520 (30 min), and HRP blocker (15 min). Slides were washed twice in 1X Wash Buffer at RT between each incubation step. After a final wash step, slides were counterstained with RNAscope Multiplex FL v2 DAPI for 30 sec and mounted with ProLong™ Gold Antifade Mountant (#P36934, Invitrogen). After drying, slides were stored in the dark at 4*°*C until imaging.

To visualize tagged transcripts, tiled images of the entirety of each RNAscope-treated section were captured using a confocal microscope (710 LSM, Zeiss) with a C-Apochromat 40x/1.2 N.A. water objective. The cortical region of each brain section was determined by comparison to the Allen Brain Atlas and outlined for analysis. Nuclei (stained with DAPI) with green staining were classified as *Sst*-expressing cells and counted using Image J Cell Counter plugin tool. *Sst*-expressing cells were counted as expressing *Htr1a* if there were at least two red puncta overlapping with the nucleus. Two transcripts were used as the threshold to reduce false positives. Percentages of *Sst*-expressing cells that contained *Htr1a* were calculated by dividing the number of cells co-expressing *Htr1a* and *Sst* by the number of *Sst*-expressing cells counted.

### Behavioral assays

All behavioral assays were conducted between 10:00 AM and 4:00 PM. To ensure the reproducibility of behavioral findings, assays were performed across multiple cohorts. In one cohort, which included 17 *Sst^Cre/+^*;*Htr1a^fl/fl^*(SST-1A cKO) and 26 sibling *Sst^+/+^*;*Htr1a^fl/fl^*control mice, the same animals underwent all behavioral assays. Both male and female mice were included, and animals were randomly assigned to experimental groups while maintaining a similar sex ratio across groups. The fear extinction test was conducted first, followed by the tail-suspension test two weeks later, and the head-twitch response test another two weeks after that. To minimize potential crosstalk between assays, at least two weeks was maintained between tests for drug washout. For other cohorts, each animal was used in only one behavioral assay.

In total, 68 mice underwent the fear extinction test, including 19 control mice treated with saline, 20 control mice treated with psilocybin, 14 SST-1A cKO mice treated with saline, and 15 SST-1A cKO mice treated with psilocybin. The tail-suspension test was performed on 84 mice, comprising 24 control mice treated with saline, 23 control mice treated with psilocybin, 18 SST-1A cKO mice treated with saline, and 19 SST-1A cKO mice treated with psilocybin. The head-twitch response test was conducted on 43 mice, including 10 control mice treated with saline, 16 control mice treated with psilocybin, 7 SST-1A cKO mice treated with saline, and 10 SST-1A cKO mice treated with psilocybin.

### Fear conditioning and extinction

The experiments were conducted using a near-infrared video fear conditioning system (MED-VFC2-SCT-M, Med Associates). Prior to each session, mice were acclimated to the behavior room for 30 min. The conditioning chambers were equipped with stainless-steel grid floors and controlled by VideoFreeze software (Med Associates). The near-infrared video camera was calibrated before testing, maintaining an average intensity of 130 au as recommended by the manufacturer. On day 1 (auditory cued fear conditioning), mice were individually placed in the chamber, which had blank straight walls and a stainless-steel grid floor, cleaned with 70% ethanol (context A). After a 3 min habituation period, each mouse received 5 presentations of an auditory tone (conditioned stimulus, CS; 4 kHz, 80 dB, 30 s duration), each of which co-terminated with a foot shock (unconditioned stimulus, US; 0.8 mA, 2 s duration). The CS-US pairings were separated by 90 s intertrial intervals. On day 3 (fear extinction), mice were administered either psilocybin (1 mg/kg, i.p.) or saline (10 mL/kg, i.p.). At 30 min post-injection, the fear extinction test commenced. The chamber was modified by inserting two black IRT acrylic sheets to create a sloped roof, and the stainless-steel grid floor was covered with a white smooth floor (context B). The chamber was cleaned with 1% acetic acid. Mice were individually placed in the chamber, given 3 min to habituate, and then exposed to 15 CS presentations without the US, with a 15 s intertrial interval. On day 4 (extinction retention), the retention procedure was conducted in context B under the same conditions as day 3, but without prior drug administration. On Day 12 (fear renewal), mice were placed in a modified conditioning chamber, characterized by a striped curved striped wall and a striped plastic floor, cleaned with Peroxigard before each animal (context C). The session followed the same habituation and CS-only exposure protocol as used on days 3 and 4. Video recordings from the near-infrared camera were analyzed using VideoFreeze software (Med Associates) with automated motion detection (detection method: linear, motion threshold: 18 au, minimum freezing duration: 30 frames = 1 s). For extinction, extinction retention, and fear renewal sessions, freezing responses were analyzed by binning the data as follows: the habituation bin included the 3 min pre-CS period, while the 15 CS presentations were showed as 5 bins by averaging freezing responses across every three consecutive tones (e.g., tones 1–3, 4–6, 7–9, 10–12, and 13–15).

### Tail suspension test

The tail-suspension test was conducted using a metal hanging apparatus constructed from optical beam parts (Thorlabs). At 24 hr prior to testing, mice received either psilocybin (1 mg/kg, i.p.) or saline (10 mL/kg, i.p.). The following day, in a tall sound-attenuating cubicle (Med Associates), each mouse was suspended by its tail using removable tape, leaving approximately 5 mm of the tail exposed. A small plastic tube was placed around the base of the tail to prevent tail climbing. The suspension lasted for 6 min, during which a video camera (acA1920, Basler) was positioned directly in front of the subject to record movement. The hanging apparatus and cubicle were thoroughly cleaned with 70% ethanol between animals. Immobility time was assessed by a blinded scorer and was defined as periods when the majority of both front and hind limbs remained motionless.

### Head-twitch response

Each mouse received an intraperitoneal injection of either psilocybin (1 mg/kg) or saline (10 mL/kg). Head-twitch responses were measured in groups of 2–3 mice, with psilocybin- and saline-treated mice typically tested simultaneously; however, each mouse was placed in a separate chamber to prevent interactions. Immediately after injection, each mouse was placed into a plexiglass chamber (4’’ × 4’’ × 4’’) with a transparent lid, positioned within a sound-attenuating cubicle (Med Associates). A high-speed video camera (acA1920, Basler) was mounted overhead to capture recordings from all chambers simultaneously. Video recordings were collected for 10 min, and chambers were thoroughly cleaned with 70% ethanol between animals. Head-twitch responses were manually scored by an experimenter blinded to the treatment conditions. While we have previously demonstrated that head twitches can be quantified using magnetic ear tags (*79*), we opted for video recording in this study to avoid potential interference of the ear tags with performance in other behavioral assays.

### Statistics

Statistical tests were conducted using Python and R studio. For electrophysiological data, mixed effects models were fit for the normalized change in firing rates (post-drug value minus pre-drug value, divided by pre-drug value) using the mixedlm function in statsmodels module in Python. We constructed a model for each cell type with fixed effects using fixed effect of treatment, along with a random effects term (intercept) for nested cell per mouse. For imaging data, mixed effects models were fit for the normalized change in calcium event rates (post-drug value minus pre-drug value, divided by pre-drug value) using the lme4 package in R. We constructed a model for each cell type using fixed effects of treatment, sex, and treatment order, along with a random effects term (intercept) for nested cell per mouse. For behavioral studies, statistical analyses were performed with GraphPad Prism 10.3. In the 4 sessions of fear extinction tests, two-factor ANOVAs were used to assess the interaction between treatment (psilocybin vs. saline) and genotype (control vs. SST-1A cKO) on the percentage of freezing across all CS presentations. Mixed-effects model with Bonferroni’s multiple comparisons test was applied to compare psilocybin:control vs. saline:control, as well as psilocybin:SST-1A cKO vs. saline:SST-1A cKO at each tone bin. For the tail-suspension test and head-twitch response, two-factor ANOVAs were used to examine the interaction between treatment (psilocybin vs. saline) and genotype (control vs. SST-1A cKO) on immobility percentage and the number of head-twitch responses. *Post hoc* comparisons with Bonferroni correction were used to assess differences between psilocybin:control and saline:control, as well as psilocybin:SST-1A cKO and saline:SST-1A cKO.

## Data and code availability

The data that support the findings and the code used to analyze the data in this study will be made publicly available at https://github.com/Kwan-Lab.

## REFERENCES

1. D. Nutt, D. Erritzoe, R. Carhart-Harris, Psychedelic Psychiatry’s Brave New World. Cell 181, 24–28 (2020).

2. F. X. Vollenweider, K. H. Preller, Psychedelic drugs: neurobiology and potential for treatment of psychiatric disorders. Nat Rev Neurosci 21, 611–624 (2020).

3. A. K. Davis et al., Effects of Psilocybin-Assisted Therapy on Major Depressive Disorder: A Randomized Clinical Trial. JAMA Psychiatry 78, 481–489 (2021).

4. R. Carhart-Harris et al., Trial of Psilocybin versus Escitalopram for Depression. N Engl J Med 384, 1402–1411 (2021).

5. G. M. Goodwin et al., Single-Dose Psilocybin for a Treatment-Resistant Episode of Major Depression. N Engl J Med 387, 1637–1648 (2022).

6. C. L. Raison et al., Single-Dose Psilocybin Treatment for Major Depressive Disorder: A Randomized Clinical Trial. JAMA 330, 843–853 (2023).

7. R. von Rotz et al., Single-dose psilocybin-assisted therapy in major depressive disorder: A placebo-controlled, double-blind, randomised clinical trial. EClinicalMedicine 56, 101809 (2023).

8. C. Liao et al., Single-nucleus transcriptomics reveals time-dependent and cell-type-specific effects of psilocybin on gene expression. biorxiv, (2025).

9. J. C. Athilingam, R. Ben-Shalom, C. M. Keeshen, V. S. Sohal, K. J. Bender, Serotonin enhances excitability and gamma frequency temporal integration in mouse prefrontal fast-spiking interneurons. Elife 6, (2017).

10. Z. Xiang, D. A. Prince, Heterogeneous actions of serotonin on interneurons in rat visual cortex. J Neurophysiol 89, 1278–1287 (2003).

11. M. V. Puig, A. Watakabe, M. Ushimaru, T. Yamamori, Y. Kawaguchi, Serotonin modulates fast-spiking interneuron and synchronous activity in the rat prefrontal cortex through 5-HT1A and 5-HT2A receptors. J Neurosci 30, 2211–2222 (2010).

12. P. Zhong, Z. Yan, Differential regulation of the excitability of prefrontal cortical fast-spiking interneurons and pyramidal neurons by serotonin and fluoxetine. PLoS One 6, e16970 (2011).

13. P. W. Sheldon, G. K. Aghajanian, Serotonin (5-HT) induces IPSPs in pyramidal layer cells of rat piriform cortex: evidence for the involvement of a 5-HT2-activated interneuron. Brain Res 506, 62–69 (1990).

14. F. Garcia-Oscos et al., Activation of 5-HT receptors inhibits GABAergic transmission by pre-and post-synaptic mechanisms in layer II/III of the juvenile rat auditory cortex. Synapse 69, 115–127 (2015).

15. J. Wood, Y. Kim, B. Moghaddam, Disruption of prefrontal cortex large scale neuronal activity by different classes of psychotomimetic drugs. J Neurosci 32, 3022–3031 (2012).

16. C. T. Golden, P. Chadderton, Psilocybin reduces low frequency oscillatory power and neuronal phase-locking in the anterior cingulate cortex of awake rodents. Sci Rep 12, 12702 (2022).

17. R. J. Olson et al., Decoupling of cortical activity from behavioral state following administration of the classic psychedelic DOI. Neuropharmacology 257, 110030 (2024).

18. J. Muir et al., Isolation of psychedelic-responsive neurons underlying anxiolytic behavioral states. Science 386, 802–810 (2024).

19. X. Xu, K. D. Roby, E. M. Callaway, Immunochemical characterization of inhibitory mouse cortical neurons: three chemically distinct classes of inhibitory cells. J Comp Neurol 518, 389–404 (2010).

20. A. Kepecs, G. Fishell, Interneuron cell types are fit to function. Nature 505, 318–326 (2014).

21. R. Tremblay, S. Lee, B. Rudy, GABAergic Interneurons in the Neocortex: From Cellular Properties to Circuits. Neuron 91, 260–292 (2016).

22. C. K. Pfeffer, M. Xue, M. He, Z. J. Huang, M. Scanziani, Inhibition of inhibition in visual cortex: the logic of connections between molecularly distinct interneurons. Nat Neurosci 16, 1068–1076 (2013).

23. L. Campagnola et al., Local connectivity and synaptic dynamics in mouse and human neocortex. Science 375, eabj5861 (2022).

24. M. Galarreta, S. Hestrin, Electrical and chemical synapses among parvalbumin fast-spiking GABAergic interneurons in adult mouse neocortex. Proc Natl Acad Sci U S A 99, 12438–12443 (2002).

25. S. Lee, I. Kruglikov, Z. J. Huang, G. Fishell, B. Rudy, A disinhibitory circuit mediates motor integration in the somatosensory cortex. Nat Neurosci 16, 1662–1670 (2013).

26. H. J. Pi et al., Cortical interneurons that specialize in disinhibitory control. Nature 503, 521–524 (2013).

27. M. Dipoppa et al., Vision and Locomotion Shape the Interactions between Neuron Types in Mouse Visual Cortex. Neuron 98, 602–615 e608 (2018).

28. D. Kvitsiani et al., Distinct behavioural and network correlates of two interneuron types in prefrontal cortex. Nature 498, 363–366 (2013).

29. L. Pinto, Y. Dan, Cell-Type-Specific Activity in Prefrontal Cortex during Goal-Directed Behavior. Neuron 87, 437–450 (2015).

30. C. Gasselin, B. Hohl, A. Vernet, S. Crochet, C. C. H. Petersen, Cell-type-specific nicotinic input disinhibits mouse barrel cortex during active sensing. Neuron 109, 778–787 e773 (2021).

31. A. Attinger, B. Wang, G. B. Keller, Visuomotor Coupling Shapes the Functional Development of Mouse Visual Cortex. Cell 169, 1291–1302 e1214 (2017).

32. J. M. Pakan et al., Behavioral-state modulation of inhibition is context-dependent and cell type specific in mouse visual cortex. Elife 5, (2016).

33. H. Taniguchi et al., A resource of Cre driver lines for genetic targeting of GABAergic neurons in cerebral cortex. Neuron 71, 995–1013 (2011).

34. L. Madisen et al., A toolbox of Cre-dependent optogenetic transgenic mice for light-induced activation and silencing. Nat Neurosci 15, 793–802 (2012).

35. J. J. Jun et al., Fully integrated silicon probes for high-density recording of neural activity. Nature 551, 232–236 (2017).

36. S. Hippenmeyer et al., A developmental switch in the response of DRG neurons to ETS transcription factor signaling. PLoS Biol 3, e159 (2005).

37. L. X. Shao et al., Psilocybin induces rapid and persistent growth of dendritic spines in frontal cortex in vivo. Neuron 109, 2535–2544 e2534 (2021).

38. Z. Yao et al., A taxonomy of transcriptomic cell types across the isocortex and hippocampal formation. Cell 184, 3222–3241.e3226 (2021).

39. N. K. Savalia, L. X. Shao, A. C. Kwan, A Dendrite-Focused Framework for Understanding the Actions of Ketamine and Psychedelics. Trends Neurosci 44, 260–275 (2021).

40. M. F. Davies, R. A. Deisz, D. A. Prince, S. J. Peroutka, Two distinct effects of 5-hydroxytryptamine on single cortical neurons. Brain Res 423, 347–352 (1987).

41. R. Araneda, R. Andrade, 5-Hydroxytryptamine2 and 5-hydroxytryptamine 1A receptors mediate opposing responses on membrane excitability in rat association cortex. Neuroscience 40, 199–412 (1991).

42. A. Fletcher et al., Electrophysiological, biochemical, neurohormonal and behavioural studies with WAY-100635, a potent, selective and silent 5-HT1A receptor antagonist. Behav Brain Res 73, 337–353 (1996).

43. B. R. Chemel, B. L. Roth, B. Armbruster, V. J. Watts, D. E. Nichols, WAY-100635 is a potent dopamine D4 receptor agonist. Psychopharmacology (Berl) 188, 244–251 (2006).

44. Y. C. Saito et al., Serotonergic Input to Orexin Neurons Plays a Role in Maintaining Wakefulness and REM Sleep Architecture. Front Neurosci 12, 892 (2018).

45. S. Maren, A. Holmes, Stress and Fear Extinction. Neuropsychopharmacology 41, 58–79 (2016).

46. G. J. Quirk, R. Garcia, F. Gonzalez-Lima, Prefrontal mechanisms in extinction of conditioned fear. Biol Psychiatry 60, 337–343 (2006).

47. S. C. Woodburn, C. M. Levitt, A. M. Koester, A. C. Kwan, Psilocybin Facilitates Fear Extinction: Importance of Dose, Context, and Serotonin Receptors. ACS Chem Neurosci 15, 3034–3043 (2024).

48. J. F. Cryan, C. Mombereau, A. Vassout, The tail suspension test as a model for assessing antidepressant activity: review of pharmacological and genetic studies in mice. Neurosci Biobehav Rev 29, 571–625 (2005).

49. T. Fuchs et al., Disinhibition of somatostatin-positive GABAergic interneurons results in an anxiolytic and antidepressant-like brain state. Mol Psychiatry 22, 920–930 (2017).

50. A. L. Halberstadt, M. Chatha, A. K. Klein, J. Wallach, S. D. Brandt, Correlation between the potency of hallucinogens in the mouse head-twitch response assay and their behavioral and subjective effects in other species. Neuropharmacology 167, 107933 (2020).

51. S. X. Chen, A. N. Kim, A. J. Peters, T. Komiyama, Subtype-specific plasticity of inhibitory circuits in motor cortex during motor learning. Nat Neurosci 18, 1109–1115 (2015).

52. A. G. Khan et al., Distinct learning-induced changes in stimulus selectivity and interactions of GABAergic interneuron classes in visual cortex. Nat Neurosci 21, 851–859 (2018).

53. J. Urban-Ciecko, A. L. Barth, Somatostatin-expressing neurons in cortical networks. Nat Rev Neurosci 17, 401–409 (2016).

54. J. J. Letzkus, S. B. Wolff, A. Luthi, Disinhibition, a Circuit Mechanism for Associative Learning and Memory. Neuron 88, 264–276 (2015).

55. E. Park, M. B. Mosso, A. L. Barth, Neocortical somatostatin neuron diversity in cognition and learning. Trends Neurosci 48, 140–155 (2025).

56. K. A. Cummings, R. L. Clem, Prefrontal somatostatin interneurons encode fear memory. Nat Neurosci 23, 61–74 (2020).

57. L. X. Shao, et al., Psilocybin’s lasting action requires pyramidal cell types and 5-HT2A receptors. Nature, in press (2025).

58. P. A. Davoudian, L. X. Shao, A. C. Kwan, Shared and Distinct Brain Regions Targeted for Immediate Early Gene Expression by Ketamine and Psilocybin. ACS Chem Neurosci 14, 468–480 (2023).

59. P. Tiwari et al., Ventral hippocampal parvalbumin interneurons gate the acute anxiolytic action of the serotonergic psychedelic DOI. Neuron 112, 3697–3714 e3696 (2024).

60. M. Luttgen, S. Ove Ogren, B. Meister, Chemical identity of 5-HT2A receptor immunoreactive neurons of the rat septal complex and dorsal hippocampus. Brain Res 1010, 156–165 (2004).

61. R. de Filippo, et al., Somatostatin interneurons activated by 5-HT(2A) receptor suppress slow oscillations in medial entorhinal cortex. Elife 10, (2021).

62. W. Munoz, R. Tremblay, D. Levenstein, B. Rudy, Layer-specific modulation of neocortical dendritic inhibition during active wakefulness. Science 355, 954–959 (2017).

63. S. J. Wu et al., Cortical somatostatin interneuron subtypes form cell-type-specific circuits. Neuron 111, 2675–2692 e2679 (2023).

64. Z. Yao et al., A high-resolution transcriptomic and spatial atlas of cell types in the whole mouse brain. Nature 624, 317–332 (2023).

65. C. A. Marcinkiewcz et al., Serotonin engages an anxiety and fear-promoting circuit in the extended amygdala. Nature 537, 97–101 (2016).

66. T. L. Anderson et al., Distinct 5-HT receptor subtypes regulate claustrum excitability by serotonin and the psychedelic, DOI. Prog Neurobiol 240, 102660 (2024).

67. H. Liu et al., Neural circuits expressing the serotonin 2C receptor regulate memory in mice and humans. Sci Adv 10, eadl2675 (2024).

68. D. Wang et al., Serotonergic afferents from the dorsal raphe decrease the excitability of pyramidal neurons in the anterior piriform cortex. Proc Natl Acad Sci U S A 117, 3239–3247 (2020).

69. I. Ferezou et al., 5-HT3 receptors mediate serotonergic fast synaptic excitation of neocortical vasoactive intestinal peptide/cholecystokinin interneurons. J Neurosci 22, 7389–7397 (2002).

70. G. K. Aghajanian, W. E. Foote, M. H. Sheard, Lysergic acid diethylamide: sensitive neuronal units in the midbrain raphe. Science 161, 706–708 (1968).

71. M. E. Trulson, B. L. Jacobs, Dissociations between the Effects of Lsd on Behavior and Raphe Unit-Activity in Freely Moving Cats. Science 205, 515–518 (1979).

72. F. Ali et al., Ketamine disinhibits dendrites and enhances calcium signals in prefrontal dendritic spines. Nat Commun 11, 72 (2020).

73. H. Homayoun, B. Moghaddam, NMDA receptor hypofunction produces opposite effects on prefrontal cortex interneurons and pyramidal neurons. J Neurosci 27, 11496–11500 (2007).

74. A. P. Buccino et al., SpikeInterface, a unified framework for spike sorting. Elife 9, (2020).

75. M. Pachitariu, N. Steinmetz, S. Kadir, M. Carandini, H. Kenneth D, Kilosort: realtime spike-sorting for extracellular electrophysiology with hundreds of channels. biorxiv, (2016).

76. P. Shamash, M. Carandini, K. D. Harris, N. A. Steinmetz, A tool for analyzing electrode tracks from slice histology. biorxiv, (2018).

77. Q. Wang et al., The Allen Mouse Brain Common Coordinate Framework: A 3D Reference Atlas. Cell 181, 936–953 e920 (2020).

78. H. Lutcke, F. Gerhard, F. Zenke, W. Gerstner, F. Helmchen, Inference of neuronal network spike dynamics and topology from calcium imaging data. Front Neural Circuits 7, 201 (2013).

79. S. J. Jefferson et al., 5-MeO-DMT modifies innate behaviors and promotes structural neural plasticity in mice. Neuropsychopharmacology 48, 1257–1266 (2023).

