## Supplementary Material for "Psilocybin reshapes cortical inhibition through selective interneuron recruitment"

**Supplemental Information**

**Figure S1. Quality metrics and waveform features of the opto-tagged PV and SST interneurons.**

**Figure S2. Effect of psilocybin on firing rates of SST interneurons by cortical region.**

**Figure S3. Effect of psilocybin on firing rates of supragranular versus infragranular SST interneurons.**

**Figure S4. Effect of psilocybin on firing rates of PV interneurons by cortical region.**

**Figure S5. Effect of psilocybin on firing rates of supragranular versus infragranular PV interneurons.**

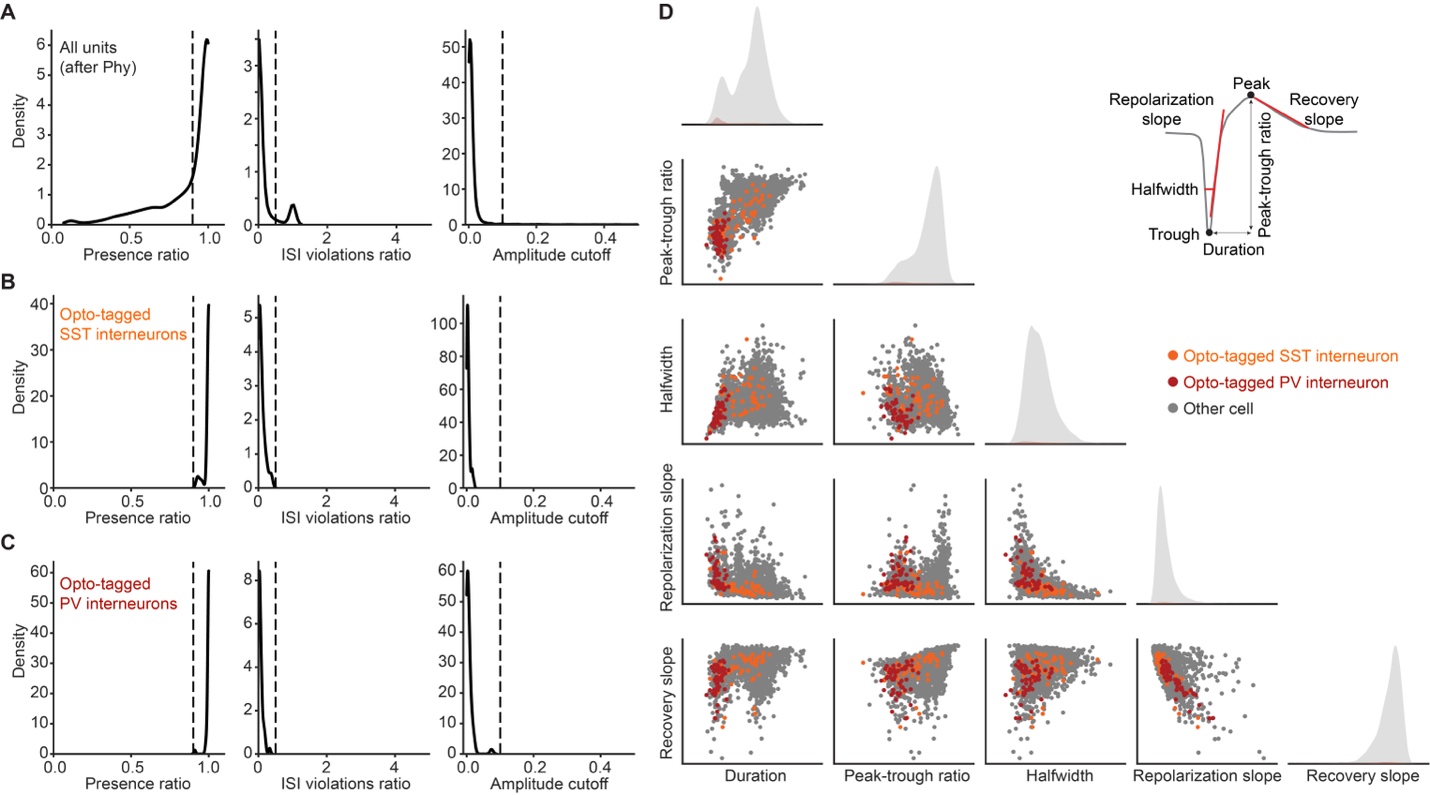

**Figure S1: Quality metrics and waveform features of the optotagged PV and SST interneurons.**

**(A)** Quality metrics for all units recorded from *Sst^Cre^*;Ai32 and *Pvalb^Cre^*;Ai32 mice after curation in Phy, but before screening for quality metrics. The curated units were included for further analysis if they met all four quality metrics: a presence ratio of ≥0.9 (dashed line), inter-spike interval violations ratio of <0.5, amplitude cutoff of <0.1 (dashed line), and isolation distance >20.

**(B)** Quality metrics for opto-tagged SST interneurons recorded from *Sst^Cre^*;Ai32 mice.

**(C)** Quality metrics for opto-tagged PV interneurons recorded from *Pvalb^Cre^*;Ai32 mice.

**(D)** Features for the mean spike waveform for opto-tagged interneurons and all other untagged cells, after screening for quality metrics, that were recorded from *Sst^Cre^*;Ai32 and *Pvalb^Cre^*;Ai32 mice.

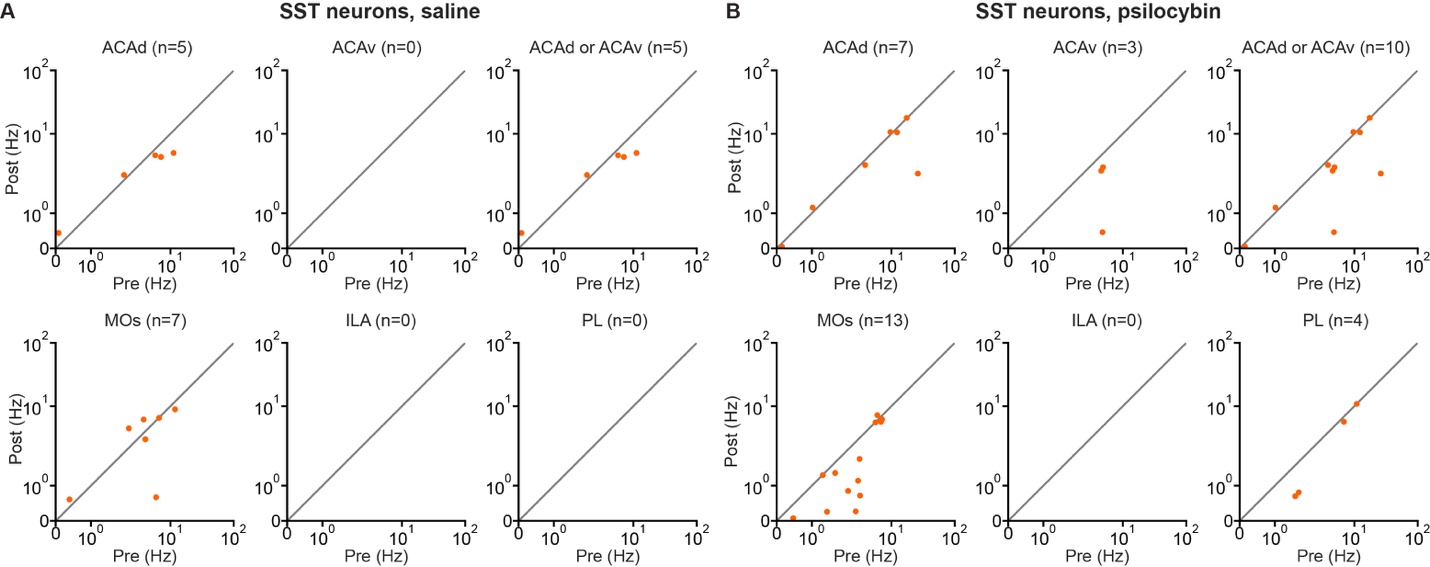

**Figure S2. Effect of psilocybin on firing rates of SST interneurons by cortical region.**

**(A)** The mean firing rates during pre- and post-saline periods for the opto-tagged SST interneurons in *Sst^Cre^*;Ai32 mice was plotted separately for units in different region (ACAd, ACAv, ACAd and ACAv, MOs, ILA, or PL). The region was estimated based on the reconstructed probe trajectory via CM-DiI fluorescence in post hoc histology. The axes use a symmetric logarithmic scale, which includes a linear region (0 to 0.03 Hz) that transitions to a logarithmic region (>0.03 Hz). Each dot represents one neuron.

**(B)** Similar to (A) for psilocybin.

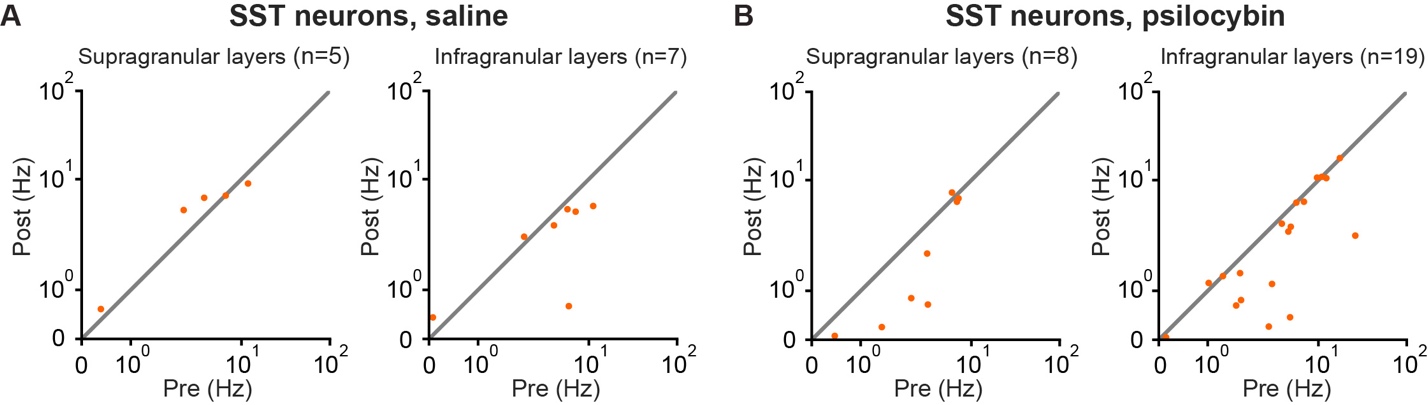

**Figure S3. Effect of psilocybin on firing rates of supragranular versus infragranular SST interneurons.**

**(A)** The mean firing rates during pre- and post-saline periods for the opto-tagged SST interneurons in *Sst^Cre^*;Ai32 mice was plotted separately for supragranular (layer 1 or layer 2/3) or infragranular (layer 5 or layer 6) units. The region was estimated based on the reconstructed probe trajectory via CM-DiI fluorescence in post hoc histology. The axes use a symmetric logarithmic scale, which includes a linear region (0 to 0.03 Hz) that transitions to a logarithmic region (>0.03 Hz). Each dot represents one neuron.

**(B)** Similar to (A) for psilocybin.

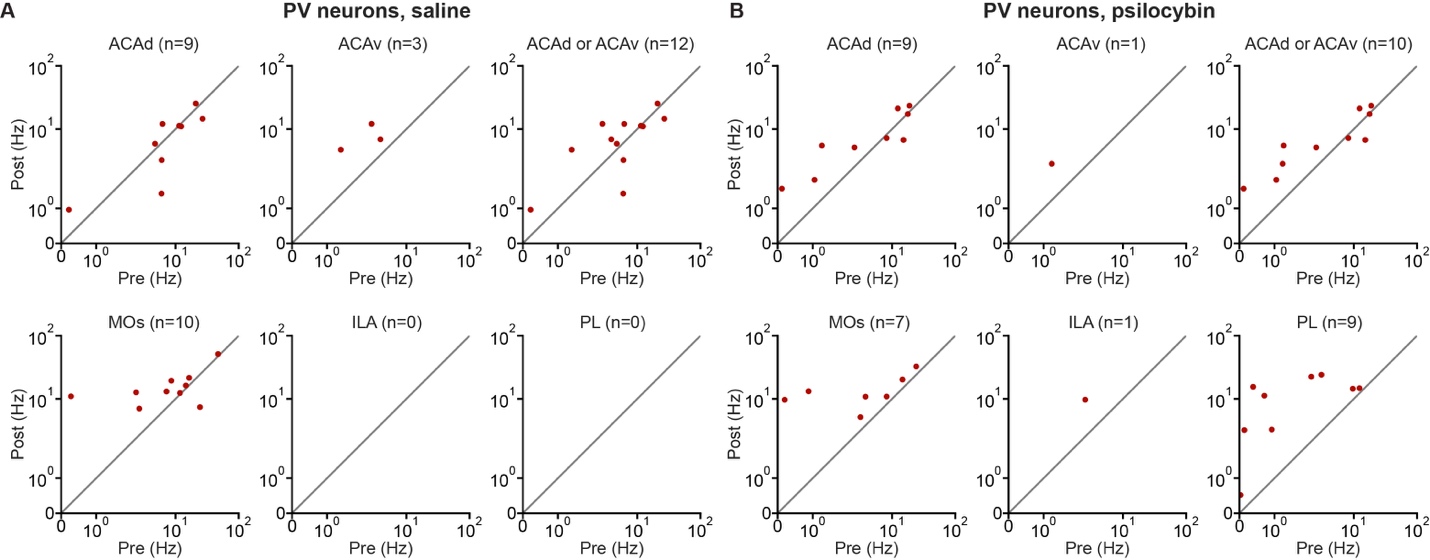

**Figure S4. Effect of psilocybin on firing rates of PV interneurons by cortical region.**

**(A)** The mean firing rates during pre- and post-saline periods for the opto-tagged PV interneurons in *Pvalb^Cre^*;Ai32 mice was plotted separately for units in different region (ACAd, ACAv, ACAd and ACAv, MOs, ILA, or PL). The region was estimated based on the reconstructed probe trajectory via CM-DiI fluorescence in post hoc histology. The axes use a symmetric logarithmic scale, which includes a linear region (0 to 0.03 Hz) that transitions to a logarithmic region (>0.03 Hz). Each dot represents one neuron.

**(B)** Similar to (A) for psilocybin.

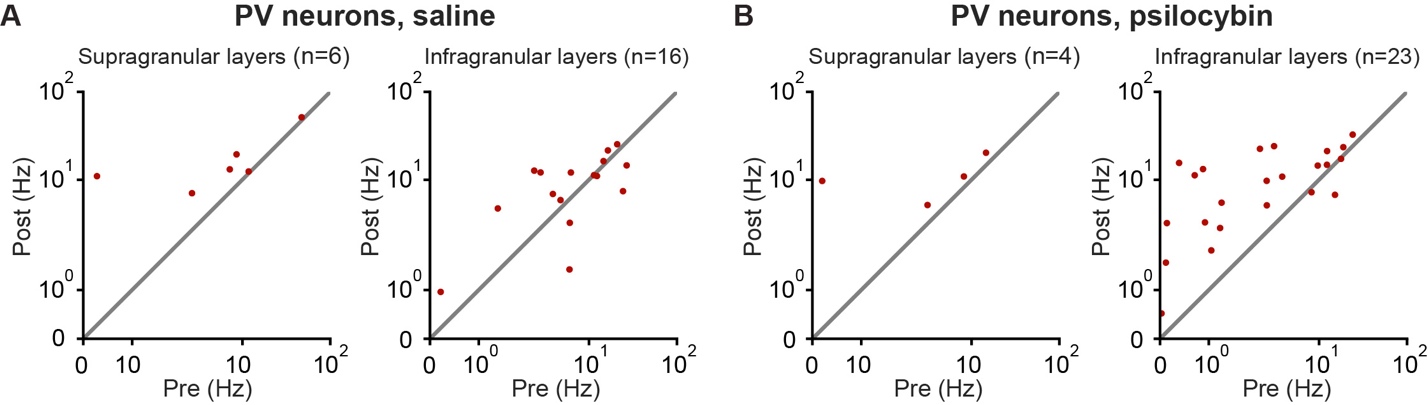

**Figure S5. Effect of psilocybin on firing rates of supragranular versus infragranular PV interneurons.**

**(A)** The mean firing rates during pre- and post-saline periods for the opto-tagged PV interneurons in *Pvalb^Cre^*;Ai32 mice was plotted separately for supragranular (layer 1 or layer 2/3) or infragranular (layer 5 or layer 6) units. The region was estimated based on the reconstructed probe trajectory via CM-DiI fluorescence in post hoc histology. The axes use a symmetric logarithmic scale, which includes a linear region (0 to 0.03 Hz) that transitions to a logarithmic region (>0.03 Hz). Each dot represents one neuron.

**(B)** Similar to (A) for psilocybin.
